# Two forms of Opa1 cooperate to complete fusion of the mitochondrial inner-membrane

**DOI:** 10.1101/739078

**Authors:** Yifan Ge, Xiaojun Shi, Sivakumar Boopathy, Julie McDonald, Adam W. Smith, Luke H. Chao

**Author notes:** Rammelkamp Center for Research and Department of Medicine, Departments of Medicine, MetroHealth System; Department of Physiology and Biophysics, School of Medicine, Case Western Reserve University, Ohio U.S.A.

## Abstract

Mitochondrial membrane dynamics is a cellular rheostat that relates metabolic function and organelle morphology. Using an *in vitro* reconstitution system, we describe a mechanism for how mitochondrial inner-membrane fusion is regulated by the ratio of two forms of Opa1. We found that the long-form of Opa1 (l-Opa1) is sufficient for membrane docking, hemifusion and low levels of content release. However, stoichiometric levels of the processed, short form of Opa1 (s-Opa1) work together with l-Opa1 to mediate efficient and fast membrane pore opening. Additionally, we found that excess levels of s-Opa1 inhibit fusion activity, as seen under conditions of altered proteostasis. These observations describe a mechanism for gating membrane fusion.

## Introduction

Mitochondrial membrane fission and fusion is essential for generating a dynamic mitochondrial network and regenerative partitioning of damaged components via mitophagy (1). Membrane rearrangement is essential for organelle function (2, 3) and contributes to diversity in mitochondrial membrane shape that can reflect metabolic and physiological specialization (4-6).

Mitochondrial membrane fusion in metazoans is catalyzed by the mitofusins (Mfn1/2) and Opa1 (the outer and inner membrane fusogens, respectively), which are members of the dynamin family of large GTPases (7, 8) (**Figure 1A**). An important series of *in vitro* studies with purified mitochondria showed that outer- and inner membrane fusion can be functionally decoupled (9, 10). Outer membrane fusion requires the Mfn1/2, while inner-membrane fusion requires Opa1. Loss of Opa1 function results in a fragmented mitochondrial network, loss of mitochondrial DNA, and loss of respiratory function (11, 12). Opa1 is the most commonly mutated gene in Dominant Optic Atrophy, a devastating pediatric condition resulting in degeneration of retinal ganglion cells. Mutations in Opa1 account for over a third of the identified cases of this form of childhood blindness (13).

**Figure 1.**
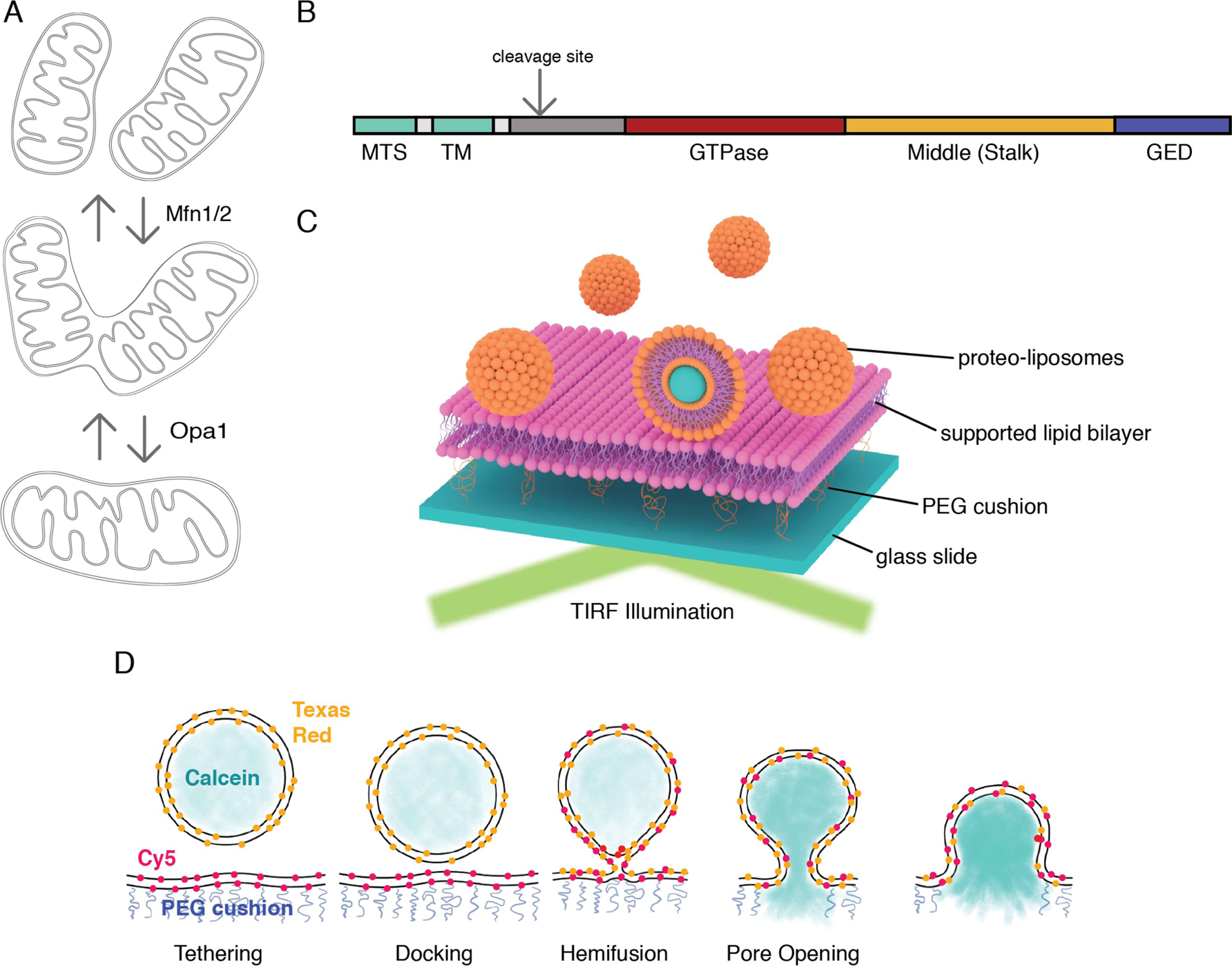
A. Mitochondrial membrane fusion involves sequential outer and inner membrane fusion. The mitofusins (Mfn1/2) catalyze outer membrane fusion. In metazoans, mitochondrial inner-membrane fusion is mediated by Opa1. B. Linear domain arrangement of l-Opa1. C. Schema of the experimental setup. D. Fusion assay. Membrane tethering, docking, lipid mixing, and content release can be distinguished using fluorescent reporters that specifically reflect each transition of the reaction.

Like dynamin, Opa1 comprises a GTPase domain, helical bundle signaling element (BSE), and stalk region (with a membrane-interaction insertion) (**Figure 1B**) (14-16). A recent crystal structure of the yeast orthologue of Opa1, Mgm1, revealed this membrane-interaction insertion is a ‘paddle’, which contains a series of hydrophobic residues that can dip into one leaflet of a membrane bilayer (17).

Opa1 is unique for a dynamin family GTPase, because it is processed to generate two forms. The unprocessed, N-terminal transmembrane anchored, long form is called l-Opa1. The proteolytically processed short form, which lacks the transmembrane anchor, is called s-Opa1 (18). Opal is processed by two proteases in a region N-terminal to the GTPase domain. Oma1 activity is stimulated by membrane depolarization (19). Yme1L activity is coupled to respiratory state. Both forms of the protein (s-Opa1 and l-Opa1) can interact with cardiolipin, a negatively charged lipid enriched in the mitochondrial inner membrane. Opa1 GTPase activity is stimulated by association with cardiolipin (20).

Recent structural studies of Mgm1 focused on a short form, s-Mgm1 construct (16). This analysis revealed a series of self-assembly interfaces in Mgm1’s stalk region. One set of interactions mediates a crystallographic dimer, and a second set, observed in both the crystal and cryo-electron tomographic (cryo-ET) reconstructions, bridge dimers in helical arrays on membrane tubes with both positive and negative curvature. The s-Mgm1 membrane tubes that formed with negative curvature are especially intriguing, because of Opa1’s recognized role in cristae regulation, and the correspondence of the *in vitro* tube topology with cristae inner-membrane invaginations (9, 21). These self-assembled states were not mediated by GTPase-domain dimers.

Integrative biophysical and structural insights have revealed how dynamin nucleotide-state is coupled to GTPase-domain dimerization, stalk-mediated self-assembly and membrane rearrangement (17, 22-24). For Opa1, the opposite reaction (fusion) is also likely to result from nucleotide-dependent conformational changes, coupled domain rearrangement, and self-assembly necessary to overcome the kinetic barriers of membrane merger. Recent crystal structure and electron cryo-tomography reconstructions reveal self-assembly interfaces, and conformational changes that rearrange cristae membranes (16). The specific fusogenic nucleotide hydrolysis-driven conformational changes remain to be distinguished.

Classic studies of Mgm1, the yeast orthologue of Opa1, show that both long and short forms are required for inner-membrane fusion (25, 26). Studies by David Chan’s group, using mammalian cells, also showed that both long and short forms of Opa1 are required (27), and that knock-down of the Opa1 processing protease Yme1L results in a more fragmented mitochondrial network (18). Since Yme1L activity is tied to respiratory state, supplying cells with substrates for oxidative phosphorylation shifts the mitochondrial network to a more tubular state. These observations led the Chan group to conclude that Opa1 processing is important for fusion. In contrast, work from the Langer group showed l-Opa1 alone was sufficient for fusion when expressed in a YME1L -/-, OMA1 -/- background (6), indicating that Opa1 processing is dispensable for fusion. Over-expression of s-Opa1 in this background resulted in mitochondrial fragmentation, which was interpreted as a result of s-Opa1 mediated fission. These directly conflicting interpretations of cellular observations have remained unreconciled. Is proteolytic processing of Opa1 required for regulating fusion, and if so, is the processing stimulatory or inhibitory?

In this study, we applied a TIRF-based supported bilayer/liposome assay (**Figure 1C**), to distinguish the sequential steps in membrane fusion that convert two apposed membranes into one continuous bilayer: tethering, membrane docking, lipid mixing (hemifusion) and content release (**Figure 1D**). This format allows control of protein levels for all components introduced into the system. Previous *in vitro* reconstitution studies from Ishihara and colleagues (28) were performed in bulk. The analysis we present here resolves individual fusion events in the TIRF field and is more sensitive than bulk measurements. In addition, our assay records kinetic data lost in ensemble averaging. Finally, the assay as applied here, can distinguish stages of fusion for individual liposomes. Tethering is observed when liposomes attach to the supported bilayer. Lipid mixing (hemifusion) is reported when a liposome dye (TexasRed) diffuses into the supported bilayer. Release of a soluble content dye (calcein) from within the liposome (loaded at quenched concentrations) indicates full pore opening. Our assay includes a content reporter dye in all conditions, so we can relate each intermediate with full fusion, for example, comparing instances where there may be lipid mixing, but no content release.

Using this *in vitro* reconstitution approach, we describe key mechanistic requirements for mitochondrial inner-membrane fusion. We report efficiency and kinetics for each step in Opa1-mediated fusion. These experiments describe the membrane activities of l-Opa1 alone, s-Opa1 alone, and l-Opa1:s-Opa1 together. We find that s-Opa1 and l-Opa1 are both required for efficient and fast pore opening, and present a mechanism for how the ratio of l-Opa1 and s-Opa1 levels regulate inner-membrane fusion. These results are compatible and expand the original yeast observations (25), explain previous cellular studies (6, 18), and contextualizes recent *in vitro* observations (28). The data presented here unambiguously describe the activities of Opa1, contributing to a more complete model for how inner-membrane fusion is regulated.

## Results

### Assay validation

We purified long and short forms of human Opa1 expressed in *Pichia pastoris*. Briefly, Opa1 was extracted from membranes using n-dodecyl-β-D-maltopyranoside (DDM) and purified by Ni-NTA and Strep-tactin affinity chromatography, and size exclusion chromatography (**Figure 2A**). A series of short isoforms are observed *in vivo* (11, 29). In this study, we focused on a short form corresponding to the S1 isoform resulting from Oma1 cleavage (**Figure 2B**). GTPase activity of purified Opa1 was confirmed by monitoring free phosphate release (**Figure 2C & D**). Opa1 activity was enhanced by the presence of cardiolipin, consistent with previous reports (**Figure 2C & D**, Figure 2-figure supplement 1) (20).

**Figure 2.**
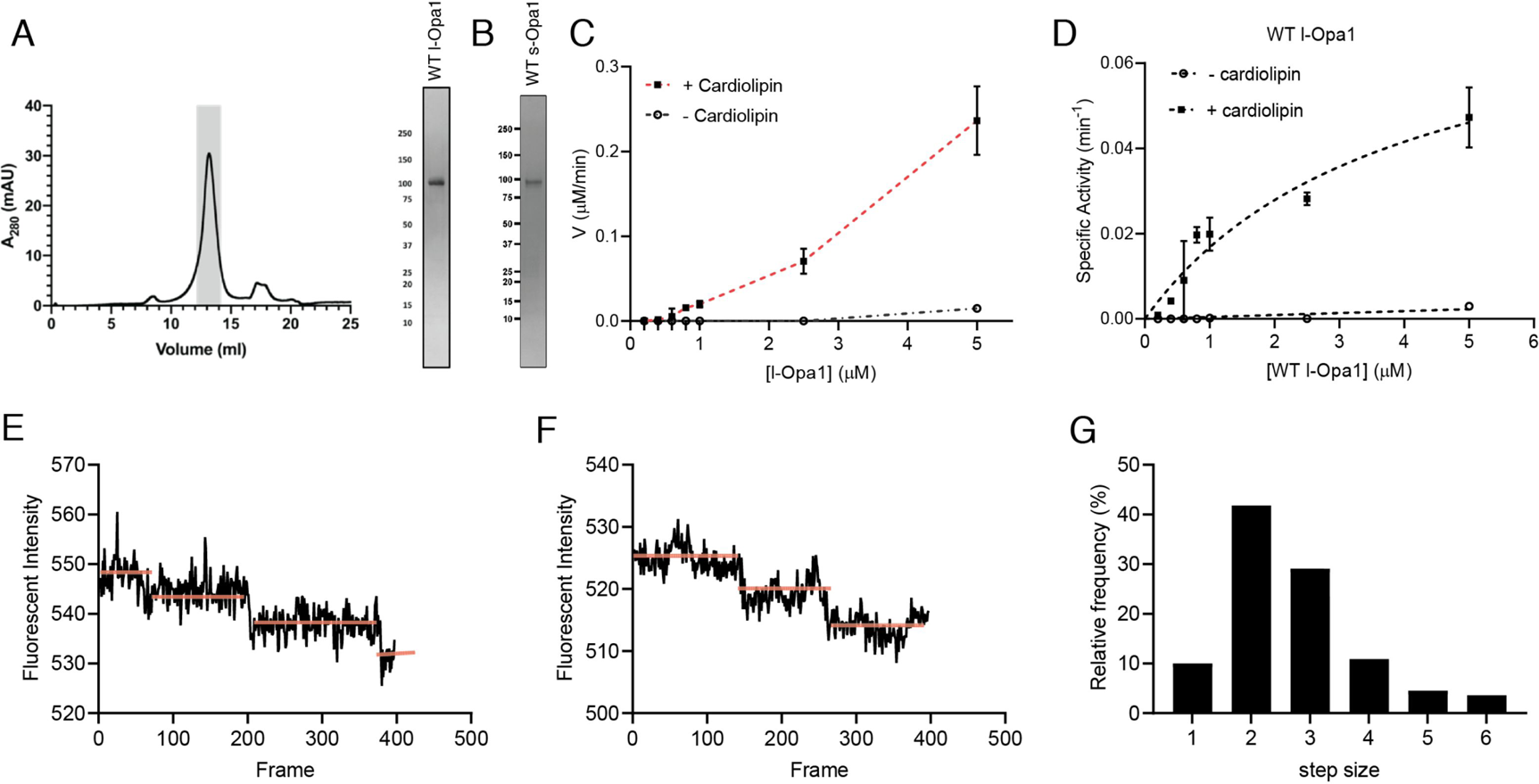
A. Representative size-exclusion chromatograph and SDS-PAGE gel of human l-Opa1 purified from *P. pastoris*. B. SDS-PAGE gel of human s-Opa1 purified from *P. pastoris*. l-Opa1 activity, with velocity (C) and specific activity (D) of GTP hydrolysis in the presence and absence of cardiolipin, while varying protein concentration of Opa1. Data are shown as mean ± SD, with error bars from 3 independent experiments. Representative single-liposome photobleaching steps (E & F) and histogram of step sizes (distribution for 110 liposomes shown) (G). Source data: Figure2-source data1.zip

We reconstituted l-Opa1 into 200 nm liposomes and supported bilayers generated by Langmuir-Blodgett/Langmuir-Schaefer methods (30). l-Opa1 was added to liposomes and a supported bilayer at an estimated protein:lipid molar ratio of 1:5000 and 1:50000, respectively. Membranes comprised DOPE (20%), Cardiolipin (20%), PI (7%), and DOPC (52.8%). Reporter dyes (e.g. Cy5-PE, TexasRed-PE) were introduced into the supported bilayer and liposome membranes, respectively, at ∼0.2 % (mol). A surfactant mixture stabilized the protein sample during incorporation. Excess detergent was removed using Bio-Beads and dialysis. We confirmed successful reconstitution by testing the stability of l-Opa1 incorporation under high salt and sodium carbonate conditions, and contrasting these results with s-Opa1 peripheral membrane association (Figure 2-figure supplement 2).

We evaluated reconstitution of l-Opa1 into both the polymer-tethered supported lipid bilayers and proteoliposomes using two approaches. First, using Fluorescence Correlation Spectroscopy (FCS), we measured the diffusion of dye-conjugated lipids and antibody-labeled protein. FCS intensity measurements confirmed ∼75% of l-Opa1 reconstituted into the bilayer in the accessible orientation. Bilayer lipid diffusion was measured as 1.46 ± 0.12 µm^2^/s, while the diffusion coefficient of bilayer-reconstituted l-Opa1 was 0.88 ± 0.10 µm^2^/s (Figure 2-figure supplement 3), which is in agreement with previous reports of lipid and reconstituted transmembrane protein diffusion (31). These measurements indicate the reconstituted l-Opa1 in the bilayer can freely diffuse, and has the potential to self-associate and oligomerize. Blue native polyacrylamide gel electrophoresis (BN-PAGE) analysis also show the potential for the purified material to self-assemble (Figure 2-figure supplement 4). FCS experiments were also performed on liposomes. FCS intensity measurements confirmed 86.7% of total introduced l-Opa1 successfully reconstituted into the liposomes. The diffusion coefficient of free antibody was 163.87 ± 22.27µm^2^/s. The diffusion coefficient for liposomes labeled with a lipid dye was 2.22 ± 0.33 µm^2^/s, and the diffusion coefficient for l-Opa1 proteoliposomes bound to a TexasRed labeled anti-His antibody was 2.12 ± 0.36 µm^2^/s (Figure 2-figure supplement 5). Second, we measured the number of l-Opa1 incorporated into liposomes by fluorescent step-bleaching of single proteoliposomes (**Figure 2E & F**). We found an average step number of 2.7 for individual l-Opa1-containing proteoliposomes labeled with TexasRed conjugated anti-His antibody, when tethered to cardiolipin containing lipid bilayers (**Figure 2G**).

### Nucleotide-dependent bilayer tethering and docking

Using the supported bilayer/liposome assay sketched in **Figure 1C**, we found that l-Opa1 tethers liposomes in a homotypic fashion (**Figure 3A**), as reported by the appearance of TexasRed puncta in the TIRF field above a l-Opa1-containing bilayer. This interaction occurred in the absence of nucleotide (apo, nucleotide-free) but was enhanced by GTP. We next investigated requirements for Opa1 tethering. In the absence of cardiolipin, addition of GTP did not change the number of tethered particles under otherwise identical conditions (**Figure 3B**). In contrast, with cardiolipin-containing liposomes and bilayers, homotypic l-Opa1:l-Opa1 tethering is enhanced by GTP. Non-hydrolyzable analogues (GMPPCP) disrupt tethering (**Figure 3C**), and a hydrolysis-dead mutant (G300E) l-Opa1 shows little tethering (Figure 3-figure supplement 1B), supporting a role for the hydrolysis transition-state in tethering, as observed for atlastin (32, 33). Bulk light scattering measurements of liposome size distributions (by NTA Nanosight) show l-Opa1-mediated liposome clustering requires the presence of GTP (Figure 3-figure supplement 2). These bulk measurements show a GTP-dependent increase in observed particle size.

**Figure 3.**
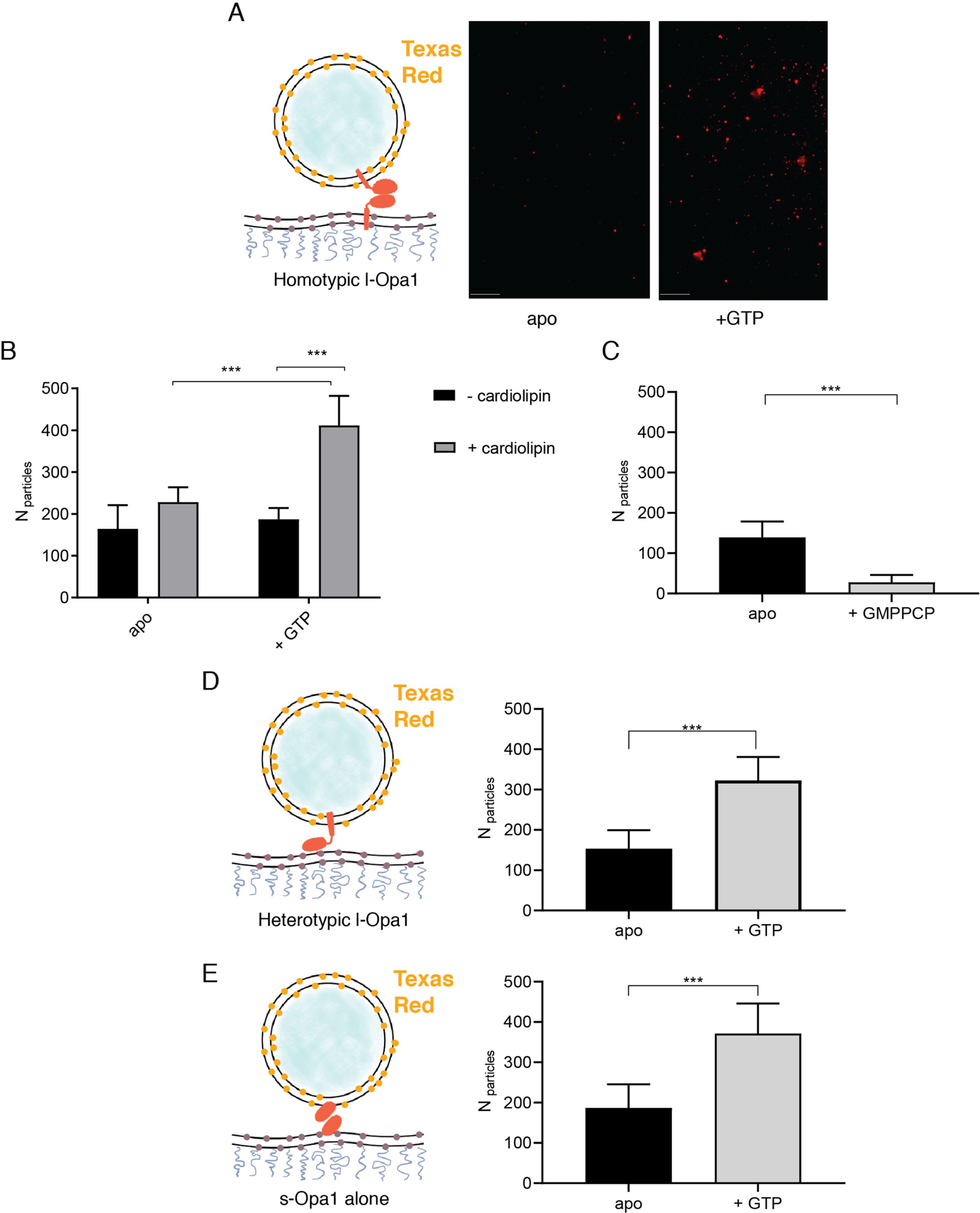
The number of liposomes tethered on the planar bilayers in a homotypic format (l-Opa1 on both bilayers) increases in the presence of GTP, when both bilayers contain cardiolipin. A. Representative images of liposomes tethered on lipid bilayer (both containing cardiolipin) before (apo, or nucleotide free) and after GTP addition. Scale bar: 5µm. B. Bar graph: In the presence of cardiolipin, addition of GTP doubles the number of liposomes. (***p<0.001, two way ANOVA). C. Addition of GMPPCP decreases amount of tethered l-Opa1 liposomes (apo, indicating nucleotide free) (P<0.005, two-way ANOVA). D. l-Opa1 in the liposome bilayer alone is sufficient to tether liposomes to a cardiolipin containing bilayer. Tethering is enhanced in the presence of GTP (apo, indicating nucleotide free) (P<0.005, two-way ANOVA). E. s-Opa1 tethers liposomes to a cardiolipin-containing bilayer. Number of tethered liposomes when both bilayer and liposomes contain 20% (mol) cardiolipin. Before addition of GTP (apo, indicating nucleotide-free), a moderate amount of liposome tethering was observed. The addition of GTP enhances this tethering effect (P<0.005, two-way ANOVA). Data are shown as mean ± SD. Error bars are from 5 independent experiments (> 10 images across one bilayer per for each experiment). Source data: Figure 3-source data1.zip

Ban, Ishihara and colleagues have observed a heterotypic, fusogenic interaction between l-Opa1 on one bilayer and cardiolipin in the opposing bilayer (28). Inspired by this work and our own observations, we considered if a heterotypic interaction between l-Opa1 and cardiolipin on the opposing membrane could contribute to l-Opa1-mediated tethering (**Figure 3D**). Indeed, we find that proteoliposomes containing l-Opa1 will tether to a cardiolipin-containing bilayer lacking any protein binding partner, presumably mediated by the lipid-binding ‘paddle’ insertion in the helical stalk region (16). This tethering is cardiolipin-dependent, as l-Opa1 containing bilayers do not tether DOPC liposomes (Figure 4-figure supplement 1B).

We next measured whether s-Opa1, lacking the transmembrane anchor, could tether membranes via membrane binding interactions that bridge the two bilayers. We observe that s-Opa1 (added at a protein:lipid molar ratio of 1:5000) can tether cardiolipin liposomes to a cardiolipin-containing planar bilayer, as observed previously for Mgm1 (34). Further, this s-Opa1 tethering is enhanced by the presence of GTP (**Figure 3E**). Previous reports observed membrane tubulation at higher concentrations of s-Opa1 (0.2 mg/ml, 1.67 nmol) (20). Under the lower s-Opa1 concentrations in our experiments (0.16 µg/ml, 2×10^-3^ nmol), the supported bilayer remains intact (before and after GTP addition), and we do not observe any evidence of tubular structures forming in the liposomes or bilayers.

Our experiments indicate that s-Opa1 alone can induce tethering. Is s-Opa1 competent for close docking of membranes? To answer this, we evaluated close bilayer approach using a reporter for when membranes are brought within FRET distances (∼40-60 Å). This FRET signal reports on close membrane docking when a TexasRed conjugated PE is within FRET distance of a Cy5-conjugated PE. We observed a low FRET signal for tethered membranes, when the FRET pair is between two supported bilayers tethered via PEG spacer (average distance between the bilayer centers of ∼7 nm), compared to a single bilayer containing both of the FRET pair (Figure 4-figure supplement 1A). When l-Opa1 is present on both bilayers (homotypic arrangement), or on only one bilayer (heterotypic arrangement), efficient docking occurs in the presence of cardiolipin, as reported by a FRET signal with efficiencies of ∼40% (**Figure 4B & C** and Figure 4-figure supplement 1A). Efficient homotypic docking requires GTP hydrolysis. GMPPCP prevents homotypic docking of l-Opa1, and abolishes the heterotypic l-Opa1 signal) (**Figure 4A**). In contrast, s-Opa1 alone does not bring the two bilayers within FRET distance, consistent with observations for s-Mgm1 tethered bilayers (**Figure 4A**) (34). The distances between two paddles in the s-Mgm1 dimer is ∼120 Å. Tethering mediated by two paddle interactions would be compatible with our observed low FRET signal when s-Opa1 engages two bilayers (17).

**Figure 4.**
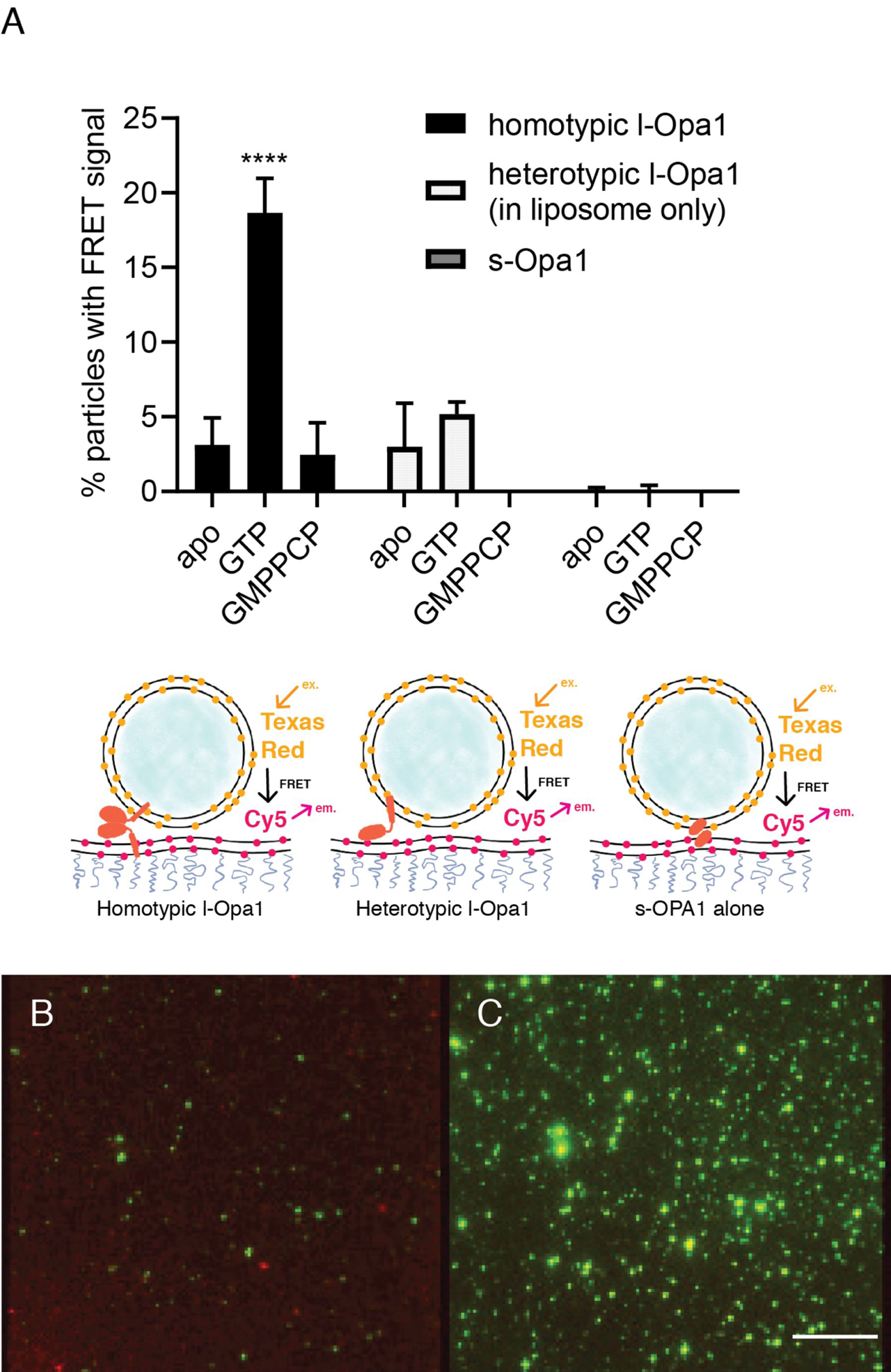
A. Homotypic l-Opa1 docks liposomes in a GTP-hydrolysis dependent manner. s-Opa1, alone is insufficient to closely dock liposomes. l-Opa1 in a heterotypic format (on the liposome alone) is competent to closely dock to a bilayer, but this docking is not stimulated by nucleotide. Bar graphs shown as mean ± SD (P<0.0001, one-way ANOVA). Error bars are from 3-5 independent experiments (each experiment with >150 particles in a given bilayer). B. In the presence of cardiolipin on both bilayers, FRET signal reports on close liposome docking mediated by l-Opa1. Left: Green = Cy5 emission signal upon excitation at 543 (TexasRed excitation). Red = Cy5 emission signal in membrane upon excitation at 633 (Cy5 excitation). Right: Green = TexasRed emission upon excitation at 543 nm (TexasRed excitation). Scale bar: 5µm. Source data: Figure 4-source data1.zip

### Hemifusion and pore opening

We find that l-Opa1, when present on only one bilayer, in a heterotypic format, can mediate close membrane docking (**Figure 4A**). To quantify hemifusion (lipid exchange), we measured the fluorescence intensity decay times for the liposome dye (TexasRed) as it diffuses into the bilayer during lipid mixing. Analysis of particle dye diffusion kinetics show that in this heterotypic format, l-Opa1 can induce hemifusion (**Figure 5A**). The hemifusion efficiency (percentage of total particles where the proteoliposome dye diffuses into the supported bilayer) for heterotypic l-Opa1 was <5% (**Figure 6A**). Previously published *in vitro* bulk liposome-based observations for heterotypic l-Opa1 lipid mixing observed hemifusion efficiencies of 5-10%, despite higher protein copy number per liposome (28). We next compared homotypic l-Opa1 catalyzed hemifusion and observed shorter mean dwell times than heterotypic l-Opa1 (**Figure 5B** & 5C, Figure 5-figure supplement 1). In our assay, we observe homotypic l-Opa1 induces hemifusion more efficiently than heterotypic l-Opa1. We measured a homotypic l-Opa1 hemifusion efficiency of ∼15% (**Figure 6A**). For comparison, *in vitro* measurements of viral membrane hemifusion, show efficiencies of ∼25-80% (35, 36). This comparison is imperfect, as viral particles have many more copies of their fusion proteins on their membrane surface and viral fusogens do not undergo multiple cycles of nucleotide hydrolysis, like Opa1.

**Figure 5.**
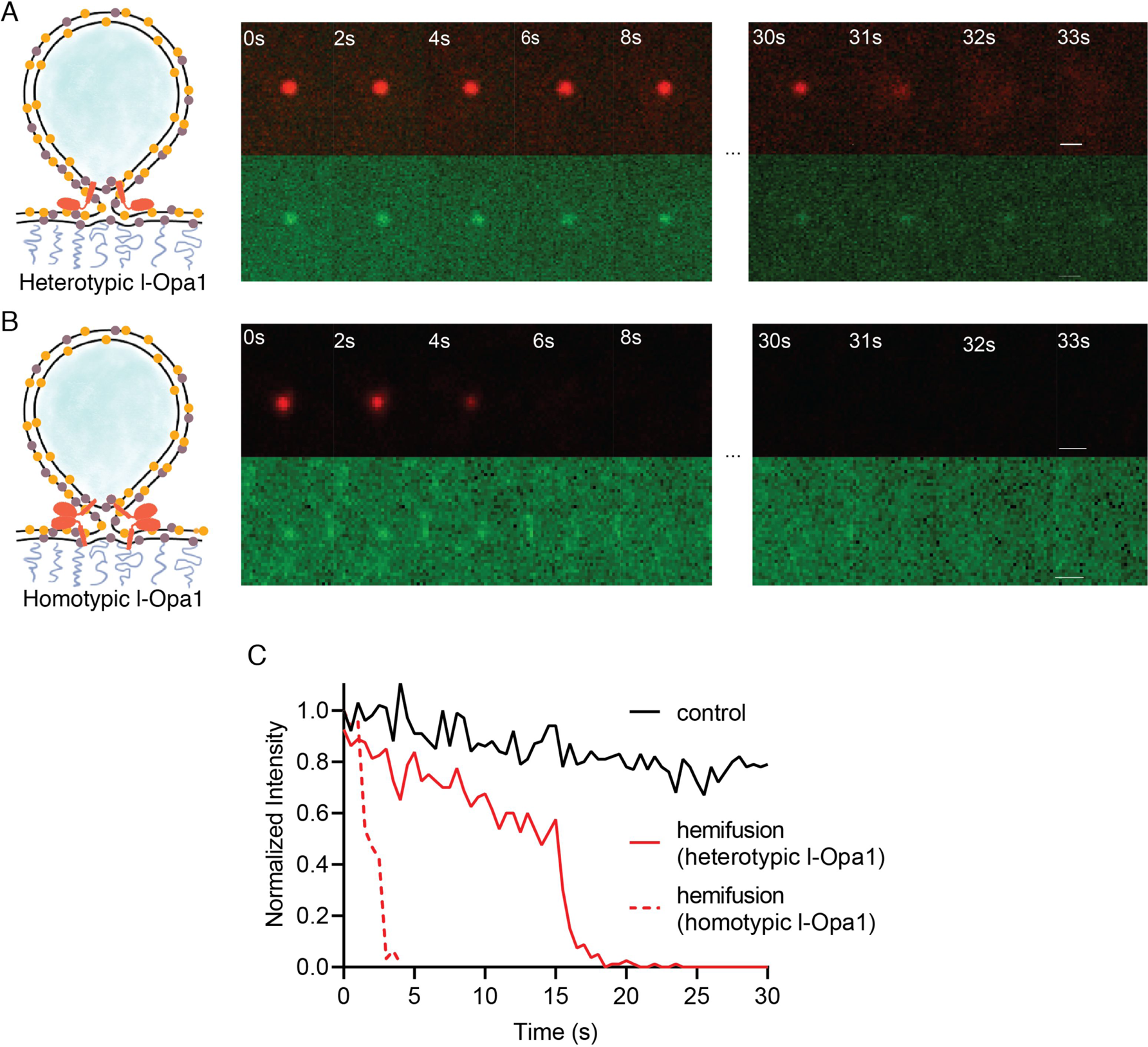
A. Heterotypic hemifusion. Top panels: time trace of proteo-liposome lipid dye diffusion (TexasRed). Bottom panels: no content release is observed for this particle (calcein signal remains quenched). Scale bar: 1 µm. B. Homotypic hemifusion. Top panels: time trace of liposome lipid dye diffusion (TexasRed). Bottom panels: no content release is observed for this particle (calcein signal remains quenched). Scale bar: 1 µm. C. Representative intensity traces of a control particle not undergoing fusion (black), with heterotypic hemifusion event (solid red), and homotypic hemifusion event (dotted red). Source data: Figure 5-source data1.zip

**Figure 6.**
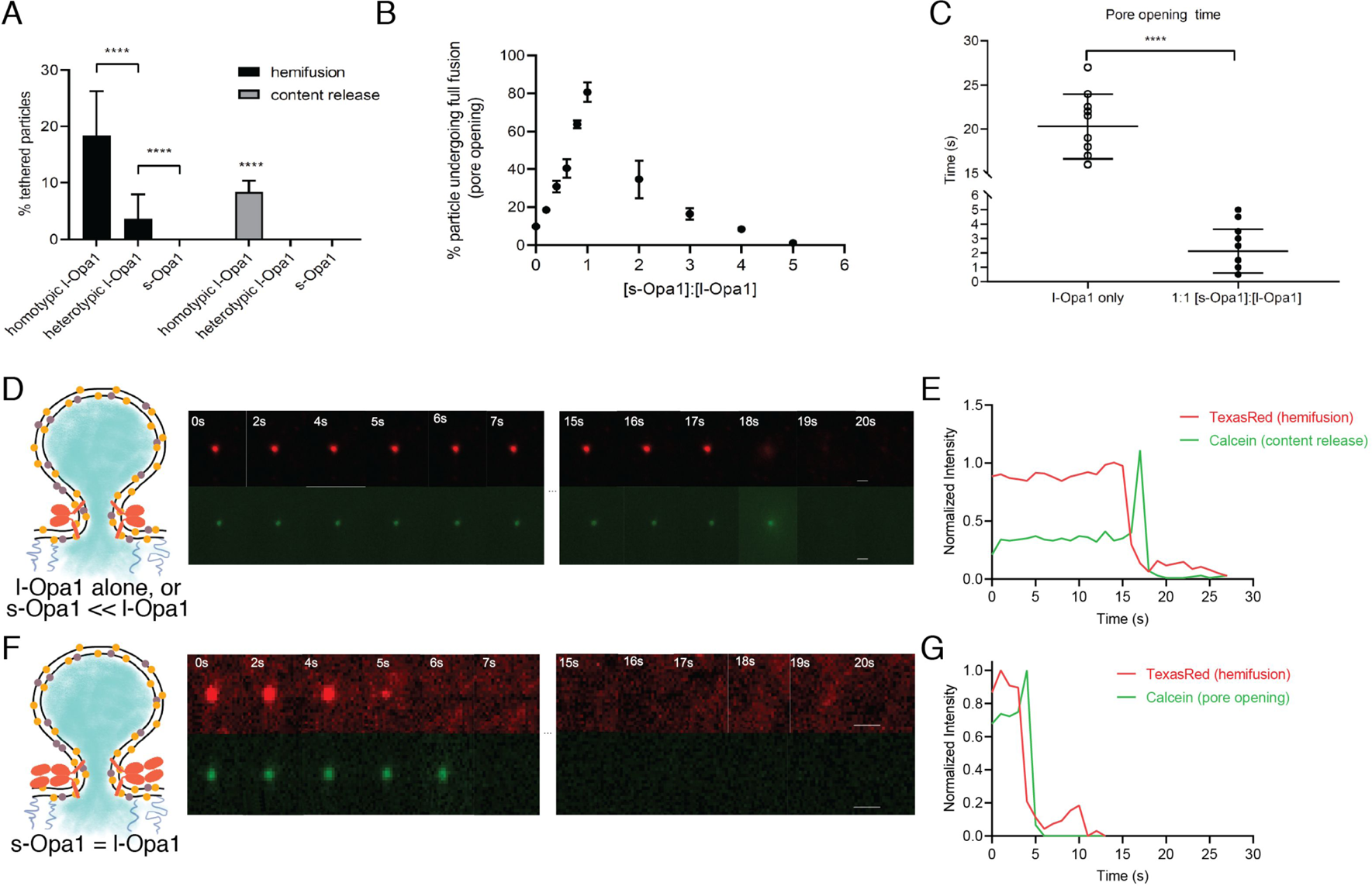
A. Hemifusion (lipid mixing) and full fusion (content release and pore opening) efficiency for homotypic l-Opa1, heterotypic l-Opa1 and s-Opa1 (P<0.001, two-way ANOVA). Bar graphs shown as mean ± SD. Error bars are from 5 different experiments (50-200 particles were analyzed per bilayer in each experiment). B. Full fusion (pore opening) efficiency at different s-Opa1:l-Opa1 ratios. Data is shown as mean ± SD. Error bars are from 4-6 experiments (80-150 particles per bilayer in each experiment). The significance of the data was confirmed using one-way ANOVA (Prism 8.3) where P<0.0001. C. Mean pore opening time in the absence of s-Opa1 and at equimolar s-Opa1. Significance of the difference was confirmed using t-test (Prism 8.3, P<0.0001). D. Representative hemifusion and pore opening fluorescence time series for homotypic l-Opa1 experiment, in the absence of s-Opa1, top and bottom panels, respectively. Scale bar: 1µm. E: representative traces of TexasRed (liposome signal) and calcein (content signal) intensity for homotypic l-Opa1 experiment. F. Representative hemifusion and pore opening fluorescence traces for a homotypic l-Opa1 experiment in the presence of equimolar s-Opa1. Scale bar: 1 µm. G: Representative trace of TexasRed (liposome signal) and calcein (content signal) intensity for a homotypic l-Opa1 experiment in the presence of equimolar s-Opa1. Source data: Figure 6-source data1.zip

Following hemifusion, pore opening is the key step where both leaflets merge and content from the two compartments can mix. We observed pore opening by monitoring content dye (calcein) release under these conditions (37). Of all homotypic tethered particles, ∼18% undergo hemifusion. Of these particles undergoing hemifusion, approximately half proceed to full fusion (8% of all homotypic tethered particles). Both s-Opa1 alone (at 0.16 µg/ml, or 2×10^-3^ nmol concentration), or l-Opa1 in the heterotypic format did not release content (**Figure 6A**). In contrast, ∼8% of homotypic l-Opa1:l-Opa1 particles undergo pore opening and content release. These observations indicate, l-Opa1 alone is sufficient for pore opening, while s-Opa1 alone or heterotypic l-Opa1 are insufficient for full fusion.

### Ratio of s-Opa1:l-Opa1 regulate pore opening efficiency and kinetics

Our observation that l-Opa1 is sufficient for pore opening leaves open the role of s-Opa1 for fusion. Previous studies suggest an active role for s-Mgm1 (the yeast orthologue of s-Opa1) in fusion (25). In this work, l-Mgm1 GTPase activity was dispensable for fusion in the presence of wild-type s-Mgm1 (25). Work in mammalian cells suggest different roles for s-Opa1. Studies from the Chan group showed Opa1 processing helps promote a tubular mitochondrial network (18). In contrast, other studies showed upregulated Opa1 processing in damaged or unhealthy mitochondria, resulting in accumulation of s-Opa1 and fragmented mitochondria (18, 28, 38). The interpretation of the latter experiments was that, in contrast to the yeast observations, s-Opa1 suppresses fusion activity. Furthermore, studies using Opa1 mutations that abolish processing of l-Opa1 to s-Opa1 suggest the short form is dispensable for fusion, and s-Opa1 may even mediate fission (39, 40). These different, and at times opposing, interpretations of experimental observations have been difficult to reconcile.

To address how s-Opa1 and l-Opa1 cooperate, we added s-Opa1 to the l-Opa1 homotypic supported bilayer/liposome fusion experiment. l-Opa1-only homotypic fusion has an average dwell time of 20 s and an efficiency of ∼10% (**Figure 6B-E** & Figure 6-figure supplement 1). Upon addition of s-Opa1, we observe a marked increase in pore opening efficiency, reaching 80% at equimolar l-Opa1 and s-Opa1 (**Figure 6B**). At equimolar levels of s-Opa1, we also observe a marked change in pore opening kinetics, with a four-fold decrease in mean dwell time (**Figure 6C**). The efficiency peaks at an equimolar ratio of s-Opa1 to l-Opa1, showing that s-Opa1 cooperates with l-Opa1 to catalyze fast and efficient fusion. When s-Opa1 levels exceed l-Opa1 (at a 2:1 ratio or greater), particles begin to detach, effectively reducing fusion efficiency. This is consistent with a dominant negative effect, where s-Opa1 likely disrupts the homotypic l-Opa1:l-Opa1 interaction. We quantified particle untethering, and observe a dose-dependent detachment of l-Opa1:l-Opa1 tethered particles with the addition of G300E s-Opa1 (**Figure S8**).

## Discussion

Our experiments suggest different assembled forms of Opa1 represent functional intermediates along the membrane fusion reaction coordinate, each of which can be a checkpoint for membrane fusion and remodeling. We show that s-Opa1 alone is sufficient to mediate membrane tethering but is unable to dock and merge lipids in the two bilayers, and thus, insufficient for hemifusion (**Figure 7A**). In contrast, l-Opa1, in a heterotypic format, can tether and hemifuse bilayers, but is unable to transition through the final step of pore opening (**Figure 7B**). Homotypic l-Opa1 can hemifuse membranes and mediate low levels of pore opening (**Figure 7C i.**). However, our results show that s-Opa1 and l-Opa1 together, synergistically catalyze efficient and fast membrane pore opening (**Figure 7C ii.**). Importantly, we find that excess levels of s-Opa1 are inhibitory for pore opening, providing a means to down-regulate fusion activity (**Figure 7C iii.**).

**Figure 7.**
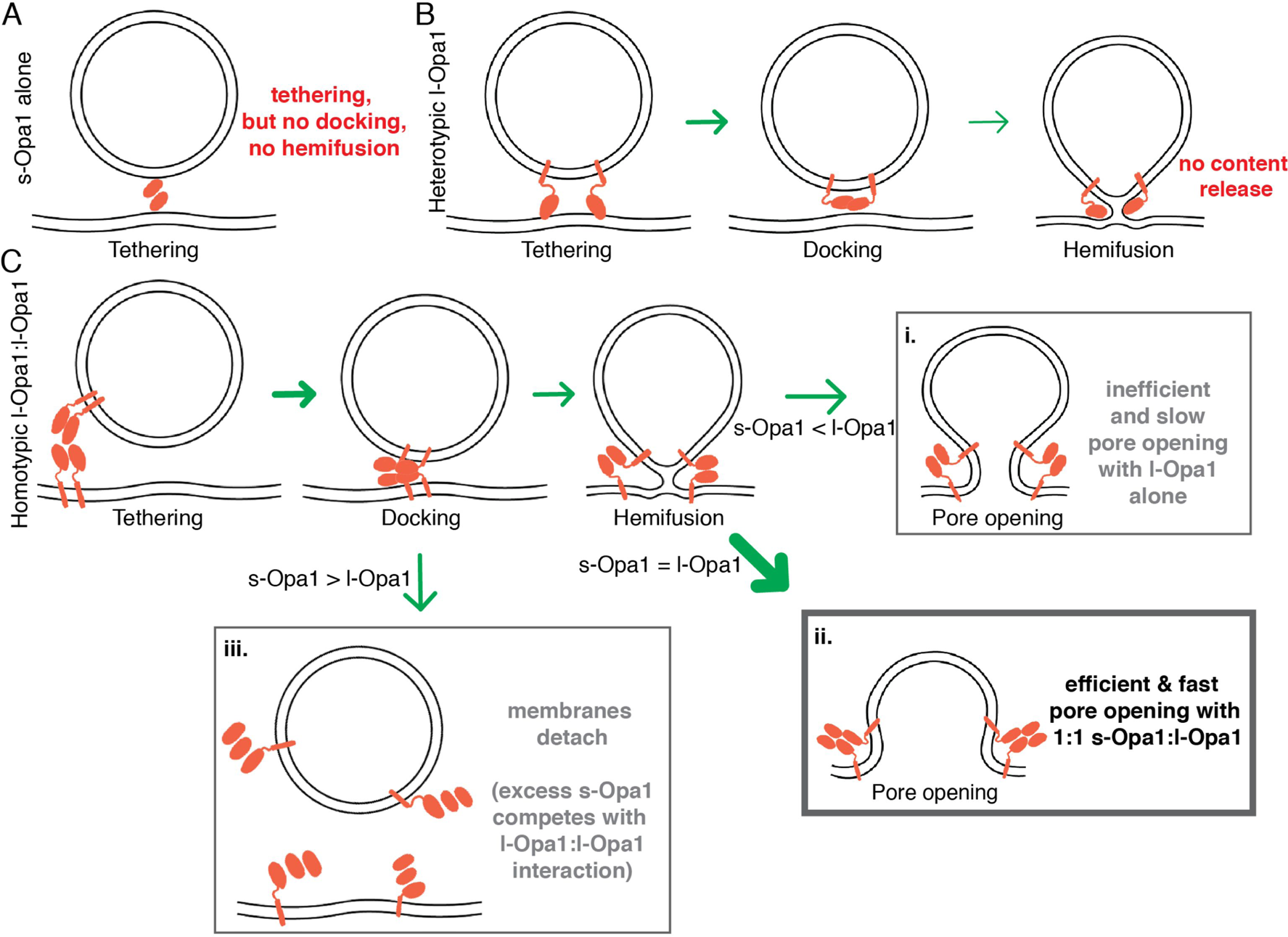
Summary model for modes of regulation in Opal-mediated membrane fusion. A. s-Opa1 alone is capable of tethering bilayers, but insufficient for close membrane docking and hemifusion. B. l-Opa1, in a heterotypic arrangement, can tether bilayers, and upon GTP stimulation promote low levels of lipid mixing, but no full fusion, pore opening or content release. C. Homotypic l-Opa1-l-Opa1 tethered bilayers can mediate full content release (i). This activity is greatly stimulated by the presence of s-Opa1, with peak activity at 1:1 s-Opa1:l-Opa1 (ii). Excess levels of s-Opa1 suppress fusion, likely through competing with the l-Opa1-l-Opa1 homotypic tethering interface (iii).

Our model proposes that l-Opa1:s-Opa1 stoichiometry, resulting from proteolytic processing, gates the final step of fusion, pore opening. Electron tomography studies of mitofusin show a unevenly distributed ring of proteins clustering at an extensive site of close membrane docking, but only local regions of pore formation (41). Our study is consistent with local regions of contact and low protein copy number mediating lipid mixing and pore formation (42). Our study would predict that s-Opa1 enrichment in regions of the mitochondrial inner-membrane would suppress fusion. This study did not explore the roles of s-Opa1 assemblies (helical or 2-dimensional) in fusion (16). Cellular validation of our proposed model, and other states, will require correlating l-Opa1:s-Opa1 ratio and protein spatial distribution with fusion efficiency and kinetics. This studied focused on the S1 form of s-Opa1. The behavior of other Opa1 splice isoforms, which vary in the processing region, remains another important area for future investigation (43).

The results and model presented here help resolve the apparent contradicting nature of the Chan and Langer cellular observations. As observed by the Langer group, l-Opa1 alone in our system, is indeed sufficient for full fusion, albeit at very low levels (6). The activity of unprocessed Opa1 was not ruled out in previous studies of Chan and colleagues (18). In contrast to the Langer group’s conclusions, we find that Opa1 processing strongly stimulates fusion activity, as observed by the Chan and colleagues (18). Under conditions of s-Opa1 overexpression, Langer *et al.* observed, a fragmented mitochondrial network. We do not see any evidence for fission or fusion, for s-Opa1 alone, under our reconstitution conditions. Instead, our data and model suggest this is due to s-Opa1 disrupting l-Opa1 activity, swinging the membrane dynamics equilibrium toward fission.

Mitochondrial dysfunction is often associated with Opa1 processing (44). The activity of the mitochondrial inner-membrane proteases, Yme1L and Oma1, is regulated by mitochondrial matrix state, thereby coupling organelle health to fusion activity (6, 40, 44–46). Basal levels of Opa1 cleavage are observed in healthy cells (18). Upon respiratory chain collapse and membrane depolarization increased protease activity elevates s-Opa1 levels, downregulating fusion (47). Our results point to the importance of basal Opa1 processing, and are consistent with observations that both over-processing and under-processing of l-Opa1 can result in a loss of function (6).

Key questions remain in understanding the function of different Opa1 conformational states, and the nature of a fusogenic Opa1 complex. Recent structural studies show s-Mgm1 self-assembles via interfaces in the stalk region (16, 48). The nucleotide-independent tethering we observe also implicate stalk region interactions, outside of a GTPase-domain dimer, in membrane tethering. How does nucleotide hydrolysis influence these interactions during fusion? Outstanding questions also remain in understanding the cooperative interplay between local membrane environment, s-Opa1, and l-Opa1. Could the cooperative activity of l-Opa1 and s-Opa1 be mediated by direct protein-protein interactions, local membrane change, or both? Could tethered states (e.g. homotypic l-Opa1 or heterotypic l-Opa1) bridge bilayers to support membrane spacings seen in cristae? Answers to these questions, and others, await further mechanistic dissection to relate protein conformational state, *in situ* architecture and physiological regulation.

## Acknowledgements

We thank members of the Chao lab for helpful discussions and review of the manuscript. LHC is grateful for support from a Charles H. Hood Foundation Child Health Research Award. We thank Fanny Ng and the Szostak Lab for technical support. Work by XS and AWS are supported by the National Science Foundation under Grant No. CHE-1753060.

## Competing interests

None

## Materials and Methods

**Table 1.**

| **Key Resources Table** |  |  |  |  |
| --- | --- | --- | --- | --- |
| **Reagent type (species) or resource** | **Designation** | **Source or reference** | **Identifiers** | **Additional information** |
| Chemical compound, drug | 18:1 (Δ9-Cis) PC (DOPC) | Avanti Polar lipids | Cat #: 850375P | |
| Chemical compound, drug | 1',3'-bis[1,2-dioleoyl-sn-glycero-3-phospho]-glycerol (sodium salt) | Avanti Polar lipids | Cat #: 710335P |  |
| Chemical compound, drug | 1,2-dioleoyl-sn-glycero-3-phosphoethanol amine-N-[methoxy(polyethylene glycol)-2000] (ammonium salt) | Avanti Polar lipids | Cat #: 880130P |  |
| Chemical compound, drug | L-α-lysophosphatidyl inositol (Liver, Bovine) (sodium salt) | Avanti Polar lipids | Cat #: 850091P | |
| Chemical compound, drug | 1-palmitoyl-2-oleoyl-sn-glycero-3-phosphoethanol amine | Avanti Polar lipids | Cat #: 850757P |  |
| Chemical compound, drug | Texas Red™<br>1,2-Dihexadecanoyl-sn-Glycero-3-Phosphoethanolamine, Triethylammonium Salt (Texas Red™ DHPE) | ThermoFisher Scientific | Cat #: T1395MP |  |
| Chemical compound, drug | 1,2-dioleoyl-sn-glycero-3-phosphoethanolamine-N-(Cyanine 5) | Avanti polar lipid | Cat #: 810335C1mg |  |
| Chemical compound, drug | Calcein | Sigma-Aldrich | Cat #: C0875; PubChem Substance ID: 24892279 |  |
| *Pichia pastoris* strain | SMD1163 (*his4, pep, prb1*) | Rapoport lab; Harvard Medical School |  |  |
| Plasmid | pPICZ A-TwinStrep-IOPA1-H<sub>10</sub> | GenScript |  | plasmid to transform and express human WT I-Opa1 (isoform1). |
| Plasmid | pPICZ A-TwinStrep-sOPA1-H<sub>10</sub> | GenScript |  | plasmid to transform and express human WT s-Opa1 (s1). |
| Plasmid | pPICZ A-TwinStrep-IOPA1 (G300E)-H<sub>10</sub> | GenScript |  | plasmid to transform and express G300E mutant of I-Opa1 (isoform 1). |
| Plasmid | pPICZ A-TwinStrep-sOPA1 (G300E)-H<sub>10</sub> | GenScript |  | plasmid to transform and express G300E mutant of s-Opa1 (s1). |
| Antibody | Anti-Opa1 antibody | NOVUS BIOLOGICALS | Cat #: NBP2-59770 | Western Blot 2 ug/ml |
| Antibody | 6x-His Tag Monoclonal Antibody (HIS.H8) | ThermoFisher Scientific | Cat #: MA1-21315 | Western Blot 1:2000 |
| Antibody | StrepMAB-Classic, HRP conjugate (2-1509-001) | IBA Lifesciences | Cat #: 2-1509-001 | Western Blot 1:2500/1:32000 |
| Antibody | Rabbit IgG HRP Linked Whole Ab | SIGMA-ALDRICH INC | Cat #: GENA934-1ML |  |
| Antibody | Mouse IgG HRP Linked Whole Ab | SIGMA-ALDRICH INC | Cat #: GENA931-1ML |  |
| Chemical compound, drug | GTP Disodium salt | SIGMA-ALDRICH INC | Cat #: 10106399001 |  |
| Chemical compound, drug | EnzChek™ Phosphate Assay Kit | ThermoFisher Scientific | Cat #: E6646 |  |
| Chemical compound, drug | GppCp (Gmppcp), Guanosine-5'-[(β,γ)-methyleno]triphosphate, Sodium salt | Jena Bioscience | Cat #: NU-402-5 |  |
| Chemical compound, drug | n-Dodecyl-β-D-Maltopyranoside | Anatrace | Cat #: D310 25 GM |  |
| Chemical compound, drug | n-Octyl-α-D-Glucopyranoside | Anatrace | Cat #: O311HA 25 GM |  |
| Chemical compound, drug | Lauryl Maltose Neopentyl Glycol (LMNG) | Anatrace | Cat #: NG310 |  |
| Chemical compound, drug | LMNG-CHS Pre-made solution | Anatrace | Cat #: NG310-CH210 |  |
| Drug | Zeocin | Invivogen | Cat #: ant-zn-1p |  |
| Resin | Ni-NTA | Biorad | Cat #: 7800812 |  |
| Resin | StrepTactin XT | IBA Lifesciences | Cat #: 2-4026-001 |  |
| Chemical compound | Biotin | IBA Lifesciences | Cat #: 2-1016-005 |  |
| Resin | Superose 6 Increase 10/300 GL | GE | Cat #: 29091596 |  |
| Reagent | TEV protease | Prepared in lab, purchased from GenScript | Cat #: Z03030 |  |
| Reagent | Benzonase Nuclease | Sigma-Aldrich | Cat #: E1014 |  |
| Reagent | Protease inhibitor cocktail | Roche | Cat #: 05056489001 |  |
| Reagent | Leupeptin | Sigma-Aldrich | Cat #: L2884 |  |
| Reagent | Pepstatin A | Sigma-Aldrich | Cat #: P5318 |  |
| Reagent | Benzamidine hydrochloride hydrate | Sigma-Aldrich | Cat #: B6506 |  |
| Reagent | Phenylmethylsulfonyl fluoride (PMSF) | Sigma-Aldrich | Cat #: 10837091001 |  |
| Reagent | Aprotinin | Sigma-Aldrich | Cat #: A1153 |  |
| Reagent | Trypsin inhibitor | Sigma-Aldrich | Cat #: T9128 |  |
| Reagent | Bestatin | GoldBio | Cat #: B-915-100 |  |
| Reagent | E-64 | GoldBio | Cat #: E-064-25 |  |
| Reagent | Phosphoramidon disodium salt | Sigma-Aldrich | Cat #: R7385 |  |
| Reagents for BN-PAGE | 3-12% Bis-Tris Protein Gels | ThermoFisher Scientific | BN1003BOX |  |
| Reagents for BN-PAGE | NativePAGE Running Buffer Kit | ThermoFisher Scientific | BN2007 |  |
| Reagents for BN-PAGE | NativePAGE 5% G-250 Sample Additive | ThermoFisher Scientific | BN2004 |  |
| Reagents for BN-PAGE | NativePAGE Sample Buffer (4X) | ThermoFisher Scientific | BN2003 |  |
| software, algorithm | Slidebook | Intelligent imaging | RRID: SCR_014300 |  |
| software,<br>algorithm | Fiji /ImageJ | Fiji | SCR_002285 |  |
| software,<br>algorithm | FCS analysis<br>tool | Smith Lab,<br>University of<br>Akron |  |  |

### Expression and purification

Genes encoding l-(residues 88-960) or s-(residues 195-960) OPA1 (UniProt O60313-1) were codon optimized for expression in *Pichia pastoris* and synthesized by GenScript (NJ, USA). The sequences encode Twin-Strep-tag, HRV 3C site, (G_4_S)_3_ linker at the N-terminus and (G_4_S)_3_ linker, TEV site, deca-histidine tag at the C-terminus. The plasmids were transformed into the methanol inducible SMD1163 strain (gift from Dr. Tom Rapoport, Harvard Medical School) and the clones exhibiting high Opa1 expression were determined using established protocols. For purification, cells expressing l-Opa1 were resuspended in buffer A (50 mM sodium phosphate, 300 mM NaCl, 1 mM 2-mercaptoethanol, pH 7.5) supplemented with benzonase nuclease and protease inhibitors and lysed using an Avestin EmulsiFlex-C50 high-pressure homogenizer. The membrane fractions were collected by ultracentrifugation at 235,000 x g for 45 min. at 4 °C. The pellet was resuspended in buffer A containing 2% DDM, (Anatrace, OH, USA) 0.1 mg/ml 18:1 cardiolipin (Avanti Polar Lipids, AL, USA) and protease inhibitors and stirred at 4 °C for 1 hr. The suspension was subjected to ultracentrifugation at 100,000 x g for 1 hr at 4 °C. The extract containing l-Opa1 was loaded onto a Ni-NTA column (Biorad, CA, USA), washed with 40 column volumes of buffer B (50 mM sodium phosphate, 350 mM NaCl, 1 mM 2-mercaptoethanol, 1 mM DDM, 0.025 mg/ml 18:1 cardiolipin, pH 7.5) containing 25 mM imidazole and 60 column volumes of buffer B containing 100 mM imidazole. The bound protein was eluted with buffer B containing 500 mM imidazole, buffer exchanged into buffer C (100 mM Tris-HCl, 150 mM NaCl, 1 mM EDTA, 1 mM 2-mercaptoethanol, 0.15 mM DDM, 0.025 mg/ml 18:1 cardiolipin, pH 8.0). In all the functional assays, the C-terminal His tag was cleaved by treatment with TEV protease and passed over the Ni-NTA and Strep-Tactin XT Superflow (IBA Life Sciences, Göttingen, Germany) columns attached in tandem. The Strep-Tactin XT column was detached, washed with buffer C and eluted with buffer C containing 50 mM biotin. The elution fractions were concentrated and subjected to size exclusion chromatography in buffer D (25 mM BIS-TRIS propane, 100 mM NaCl, 1 mM TCEP, 0.025 mg/ml 18:1 cardiolipin, pH 7.5, 0.01% LMNG, 0.001% CHS). s-OPA1 was purified using a similar approach but with one difference: post lysis, the DDM was added to the unclarified lysate at 0.5% concentration and stirred for 30 min. – 1 hr. at 4 °C prior to ultracentrifugation. The supernatant was applied directly to the Ni-NTA column.

### GTPase activity assay

The GTPase activity of purified Opa1 was analyzed using EnzCheck Phosphate Assay Kit (Thermo Fisher, USA) according to the vendor’s protocol. Each condition was performed in triplicate. The GTPase assay buffers contained 25 mM HEPES, 60 mM NaCl, 100 mM KCl, 0.5 mM MgCl_2_ with 0.15 mM DDM. 60 µM GTP was added immediately before data collection. To compare the effect of cardiolipin on GTPase activity, additional 0.5 mg/ml Cardiolipin was dissolved in the reaction buffer and added to the reaction to a final concentration of 0.02 mg/ml. The absorbance at 340 nm of each reaction mixture was recorded using SpectraMax i3 plate reader (Molecular Devices) every 30 seconds. Experiments were performed in triplicate. Resulting Pi concentration was fitted to a single-phase exponential-decay, specific activity data were fitted to a Michaelis-Menten equation (GraphPad Prism 8.1).

### Preparation of polymer-tethered lipid bilayers

Lipid reagents, including 1,2-dioleoyl-sn-glycero-3-phosphocholine, (DOPC); 1,2-dioleoyl-sn-glycero-3-phosphoethanolamine-N-[methoxy(polyethylene glycol)-2000] (DOPE-PEG2000), L-α-phosphatidylinositol (Liver PI) and 1’,3’-bis[1,2-dioleoyl-sn-glycero-3-phospho]-glycerol (cardiolipin) were purchased from Avanti Polar Lipids (AL, USA). To fabricate the polymer-tethered lipid bilayers, we combined Langmuir-Blodgett and Langmuir-Schaefer techniques, using a Langmuir-Blodgett Trough (KSV-NIMA, NY, USA) (31, 49). For cardiolipin-free lipid bilayers, a lipid mixture with DOPC with 5 % (mol) DOPE-PEG2000 and 0.2 % (mol) Cy5-DSPE at the total concentration of 1 mg/ml was spread on the air water interface in a Langmuir trough. The surface pressure was kept at 30 mN/m for 30 minutes before dipping. The first lipid monolayer was transferred to the glass substrate (25 mm diameter glass cover slide, Fisher Scientific, USA) through Langmuir-Blodgett dipping, where the dipper was moved up at a speed of 22.5 mm/min. The second leaflet of the bilayer was assembled through Langmuir-Schaefer transfer after 1 mg/ml of DOPC with 0.2 % (mol) Cy5-PE (Avanti Polar Lipids, AL, USA) was applied to an air-water interface and kept at a surface pressure of 30 mN/m.

Lipid bilayer with cardiolipin was fabricated in a similar manner, where the bottom leaflet included 7 % (mol) Liver PI, 20 % (mol) cardiolipin, 20 % (mol) DOPE, 5 % (mol) DOPE-PEG2000, 0.2% (mol) Cy5-PE and 47.8% DOPC. The composition of the top leaflet of the bilayer was identical except for the absence of DOPE-PEG2000. To match the area/molecule at the air-water interface between CL-free and CL-containing bilayer, the film pressure was kept at 37 mN/m. The final average area per lipid, which is the key factor affecting lipid lateral mobility, was kept constant at a A_lipid_ = 65 Å^2^ (50).

Double bilayers were fabricated according to previous reports (51). The first bilayer containing DOPC with 5 % (mol) DSPE-PEG2000-Maleimide (Avanti Polar Lipids, AL, USA) and 0.2 % (mol) Cy5-DOPE in both inner and outer leaflets was made using Langmuir-Blodgett/Langmuir-Schaefer methods. The second planar lipid bilayer was formed by fusion of lipid vesicles and removal of non-fused vesicles. Lipid vesicles were formed by hydrating dried lipid films with DOPC, 0.2 % (mol) TexasRed-DHPE and 5 % (mol) of linker lipid (DPTE, AL, USA) in a 0.1 mM sucrose/1 mM CaCl_2_ solution. The lipid suspension was heated for 1.5 hours at 75 °C, and added to the first bilayer in a 0.1 mM glucose/1 mM CaCl_2_ solution. After 2 hours of incubation, additional vesicles were removed by extensive rinsing. The bilayer was then imaged by TIRF microscope.

### Reconstitution of l-Opa1 into lipid bilayers

Purified l-Opa1 was first desalted into 25 mM Bis-Tris buffer with 150mM NaCl containing 1.2 nM DDM and 0.4 µg/L of cardiolipin to remove extra surfactant during purification. The resulting protein was added to each bilayer to the total amount of 1.3×10^-12^ mol (protein:lipid 1:10000) together with a surfactant mixture of 1.2 nM of DDM and 1.1 nM n-Octyl-β-D-Glucopyranoside (OG, Anatrace, OH, USA). The protein was incubated for 2 hours before removal of the surfactant. To remove the surfactant, Bio-Beads SM2 (Bio-Rad, CA, USA) was added to the solution at a final concentration of 10 µg beads per mL of solution and incubated for 10 minutes. Buffer with 25 mM Bis-Tris and 150 mM NaCl was applied to remove the Bio-beads with extensive washing. Successful reconstitution was determined using fluorescent correlation spectroscopy assay as described in the supplemental materials.

### Preparation of liposomes and proteoliposomes

To prepare calcein (MilliporeSigma, MA, USA) encapsulated liposomes, lipid mixtures (7 % (mol) PI, 20% cardiolipin, 20% PE, 0.2% TexasRed-PE, DOPC (52.8%)), were dissolved in chloroform and dried under argon flow for 25 minutes. The resulting lipid membrane was mixed in 25 mM Bis-Tris with 150mM NaCl and 50 mM calcein through vigorous vortexing. Lipid membranes were further hydrated by incubating the mixtures under 70 °C for 30 min. Large unilamellar vesicles (LUVs) were prepared by extrusion (15 to 20 times) using a mini-extruder with 200 nm polycarbonate membrane.

Proteoliposomes were prepared by adding purified l-Opa1 in 0.1 µM DDM to prepared liposomes at a protein: lipid of 1:5000 (2.5 µg l-Opa1 for 0.2 mg liposome) and incubated for 2 hours. Surfactant was removed by dialysis overnight under 4 °C using a 3.5 KDa dialysis cassette. Excess calcein was removed using a PD-10 desalting column. The final concentration of liposome was determined by TexasRed absorbance, measured in a SpectraMax i3 plate reader (Molecular Devices).

To evaluate l-Opa1 reconstitution into proteoliposomes, dye free liposome was prepared with TexasRed conjugated anti-His tag Antibody (ThermoFisher) by mixing lipids with antibody containing buffer. TexasRed Labeling efficiency of the antibody was calculated to be 1.05 according to the vendor’s protocol. Antibodies were added at a concentration of 2.6 µg/ml to 0.2 mg/ml liposome. Following hydration through vortexing at room temperature for 15 minutes, Large unilamellar vesicles were formed following 20 times extrusion procedure described above. Liposomes labeled with 0.02 % (mol) TexasRed-PE were also prepared as a standard for quantifying reconstitution rate.

For the co-flotation analysis, 200 µl of 20 mg/ml TexasRed-DHPE (0.2 % (mol)) labeled proteoliposome (reconstitution ratio, protein:lipid 1:5000) was loaded to sucrose gradient (with steps of 0%, 15%, 30%, 60%). The volume of each fraction was 800 µl. Sucrose solutions were prepared in Bis-Tris buffer (25mM Bis-Tris, 150 mM NaCl, pH 7.4). Samples were then centrifuged using a high-speed centrifuge equipped with SW 55i rotor (Beckmann Coulter, CA, USA) for 2.5 hrs at a speed of 30000 xg. For high salt and carbonate treatment, the same amount of proteoliposome was redistributed in Bis-Tris buffer with 500 mM NaCl (pH 7.4) and buffer containing 50 mM Na_2_CO_3_ and 50 mM NaCl (pH 8.2), respectively. The resulting suspension was loaded in gradient for separation. After centrifugation, all fractions were collected and concentrated to 40 µl. Fractions were detected by western blot and then analyzed by ImageJ. The presence of liposomes was detected by absorbance at 590 nm using a DeNovix FX photometer (DeNovix, Inc).

### Fluorescent Correlation Spectroscopy

Fluorescence correlation spectroscopy (FCS) was performed using a home-built PIE-FCCS system (52, 53). Two pulsed laser beams with wavelengths of 488 nm (9.7 MHz, 5 ps) and 561 nm (9.7 MHz, 5 ps) were filtered out from a supercontinuum white light fiber laser (SuperK NKT Photonics, Birkerod, Denmark) and used as excitation beams. The laser beams were sent through a 100X TIRF objective (NA 1.47, oil, Nikon Corp., Tokyo, Japan) to excite the samples in solution or on bilayer. The emission photons were guided through a common 50 μm diameter pinhole. The light was spectrally separated by a 560 nm high-pass filter (AC254-100-A-ML, Thorlabs), further filtered by respective bandpass filters (green, 520/44 nm [FF01-520/44-25]; red, 612/69 nm [FF01-621/69-25], Semrock), and finally reach two single photon avalanche diode (SPAD) detectors (Micro Photon Devices). The synchronized photon data was collected using a time correlated single photon counting (TCSPC) module (PicoHarp 300, PicoQuant, Berlin, Germany).

The collected photon data was transformed into correlation functions with a home written MATLAB code. The correlation functions were fitted using two-dimensional (1) or three-dimensional (2) Brownian diffusion model for bilayer or solution samples respectively.

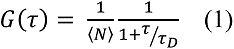

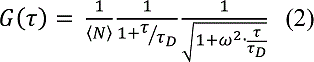

Where N is the average number of particles in the system, ω is the waist of the excitation beam, and τD is the dwell time that can be used to calculate the diffusion coefficient (D) of the particles. (52)

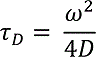

Measurements were made on buffers with evenly distributed liposomes, proteoliposomes and antibodies in a glass-bottom 96 well plate at room temperature. The plates were pre-coated with lipid bilayer fabricated from 100 nm DOPC liposomes. For each solution, data was collected in five successive 15 second increments.

For characterization of l-Opa1 reconstitution into planar bilayers, an anti-Opa1 C-terminal antibody (Novus Biologicals, CO, USA) was used. The antibody was labeled by TexasRed (Texas Red™-X Protein Labeling Kit, ThermoFisher, CA, USA). Labeling efficiency of the antibody was determined as 1.52 TexasRed/antibody, as determined by NanoDrop (ThermoFisher, CA, USA). The labeled antibody was added to l-Opa1 in the supported bilayer at twice the total introduced Opa1 concentration. Excess antibody was removed by extensive rinsing.

To estimate reconstitution efficiency, 0.002 % (mol) l-Opa1 was added to the bilayer. In a separate experiment 0.002 % (mol) TexasRed-PE was introduced to the bilayer. The reconstitution efficiency was calculated from the anti-l-Opa1 antibody TexasRed fluorophore density divided by the TexasRed-PE fluorphore density, normalized by the antibody labeling efficiency (1.5 dye molecules/antibody).

### Total Internal Reflection Fluorescent Microscopy (TIRF)

Liposome docking and lipid exchange events were imaged using a Vector TIRF system (Intelligent Imaging Innovations, Inc, Denver, CO, USA) equipped with a W-view Gemini system (Hamamatsu photonics, Bridgewater, NJ). TIRF images were acquired using a 100X oil immersion objective (Ziess, N.A 1.4). A 543 nm laser was used for the analysis of TexasRed-PE embedded liposomes and proteoliposomes, while a 633 nm laser was applied for the analysis of Cy5-PE embedded in the planar lipid bilayer. Fluorescent emission was simultaneously observed through a 609-emission filter with a band width of 40 nm and a 698-emission filter with a band width of 70 nm. The microscope system was equipped with a Prime 95B scientific CMOS camera (Photometrics), maintained at −10 °C. Images were taken at room temperature, before adding any liposome or proteoliposome, after 15 mins of addition, and after 30 mins of adding GTP (1 mM) and MgCl_2_ (1 mM). Each data point was acquired from 5 different bilayers, each bilayer data contains 5-10 particles on average.

Dwell times for hemifused particles were recorded from the moment of GTP addition for pre-tethered particles, until the time of half-maximal TexasRed signal decay. Full fusion events were recorded by monitoring the calcein channel at particle locations identified through the TexasRed signal. Particle identification and localization used both uTrack(54) and Slidebook (Intelligent Imaging Innovations, Inc., Denver, CO) built-in algorithms. To calibrate the point spread function 100 nm and 50 nm fluorescent particles (ThermoFisher Scientific) were used. 2D Gaussian detection were applied in both cases. 2-way ANOVA tests was done using GraphPad Prism. Intensity and distribution of the particles were analyzed using ImageJ.

For analysis of protein reconstitution in proteoliposome (stoichiometry), a TIRF microscope modified from an inverted microscope (Nikon Eclipse Ti, Nikon Instruments) was used. A 561 nm diode laser (OBIS, Coherent Inc., Santa Clara, USA) was applied at TIRF angle through a 100X TIRF objective (NA 1.47, oil, Nikon) and the fluorescence signals were collected by an EMCCD camera (Evolve 512, Photometrics).

### Nanosight NTA analysis

A NTA300 Nanosight instrument was used to evaluate size distribution of liposome and proteoliposome under different conditions. The equipment was equipped with a 405 nm laser and a CMOS camera. 1 ml of 0.1 µg/ml sample was measured, to reach the recommended particle number of 1× 10^8^ particles/mL (corresponding to the dilution factor of 1:100,000). Image acquisition were conducted for 40 sec for each acquisition and repeated for 10 times for every injection. Three parallel samples were examined for the determination of size distribution. Under each run, the camera level was set to 12 and the detection threshold was set at 3.

### Blue native polyacrylamide gel electrophoresis (BN-PAGE)

Bis-Tris gradient gels (3-12%) were purchased from ThermoFisher Scientific (Cat. No. BN1003BOX) and BN-PAGE was performed according to manufacturer’s instructions. Gel samples (10 μl) were prepared by mixing indicated quantity of Opa1 with sample buffer containing 0.25% Coomassie G-250 and 1 mM DDM. For experiments involving l-Opa1 and s-Opa1 mixtures, the samples were incubated on ice for 10 min before loading. The cathode buffer contained 1 mM DDM and electrophoresis was performed at 4°C with an ice jacket surrounding the apparatus.

---

**Figure 2 – figure supplement 1.**
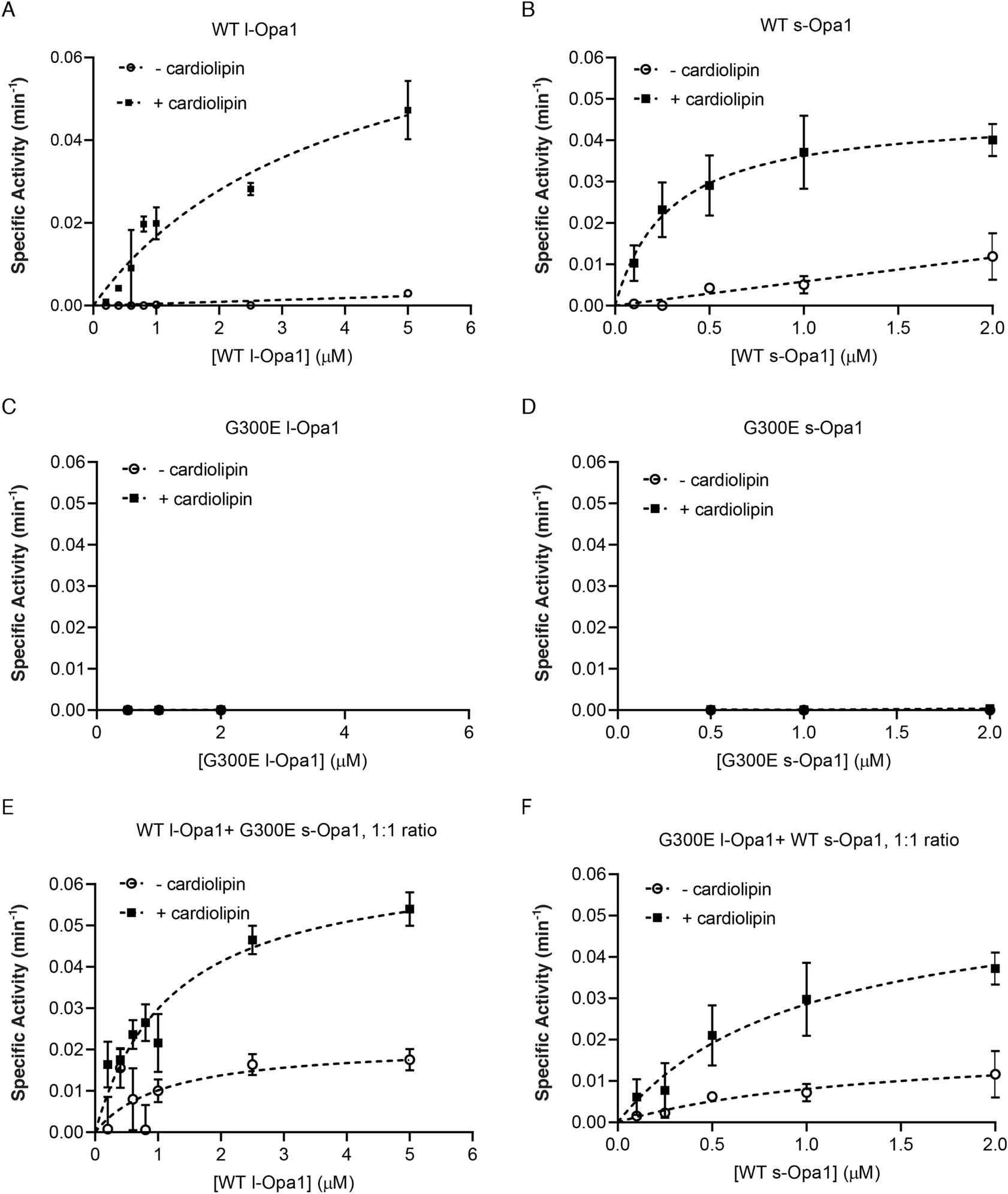
GTP hydrolysis (GTPase) activity of l-Opa1 (A) and s-Opa1 (B) in the presence and absence of cardiolipin. Both G300E l-Opa1 and G300E s-Opa1 do not show any GTPase activity (C & D). Mixing G300E s-Opa1 with WT l-Opa1 at 1:1 molar ratio (E) does not alter the GTPase activity of, detergent solubilized, WT l-Opa1 significantly (E and A, P>0.2, t-test). A similar effect is seen upon addition of G300E l-Opa1 to WT s-Opa1 at 1:1 ratio (F). Under these conditions, s-Opa1 GTPase activity is similar to s-Opa1 alone (F & B, P>0.2, t-test). Data shown as mean ± SD, error bars from 3 experiments. Source data: Figure 2-fig sup 1-source data1.zip

**Figure 2 – figure supplement 2.**
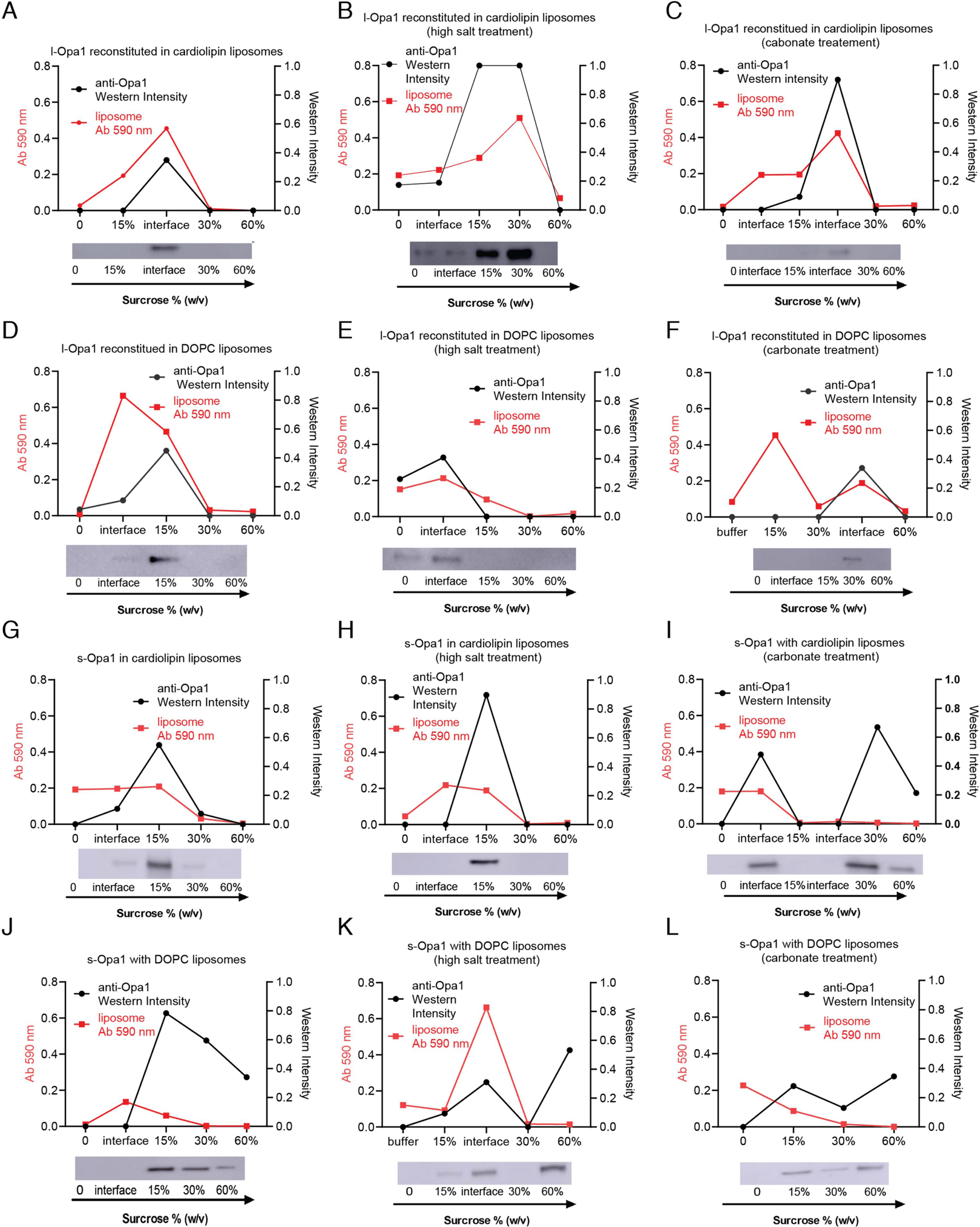
Liposome co-flotation analysis: Reconstituted l-Opa1 co-floats with liposomes both with and without cardiolipin (A & D). Liposomes were labeled with 0.2 % (mol) TexasRed-DHPE and their distribution was confirmed by liposome dye absorbance at 590 nm. Opa1 distribution was analyzed by Western blot. Opa1/liposome fractions was mostly found near 15∼30% sucrose. This reconstitution is stable under high salt (B & E) or carbonate conditions (C & F). s-Opa1 interacts with liposomes in a cardiolipin-dependent manner (G-L). This interaction is resistant to high salt (H) but sensitive to carbonate treatment (I), where the protein was found in the bottom fractions lacking liposome (60% sucrose). s-Opa1 does not associate with DOPC liposomes (J-L). These results indicate that l-Opa1 was successfully reconstituted through integral transmembrane region, whereas the s-Opa1 bilayer-association is through a cardiolipin:s-Opa1 peripheral membrane interaction. Source data: Figure 2-fig sup 2-source data1.zip

**Figure 2 – figure supplement 3.**
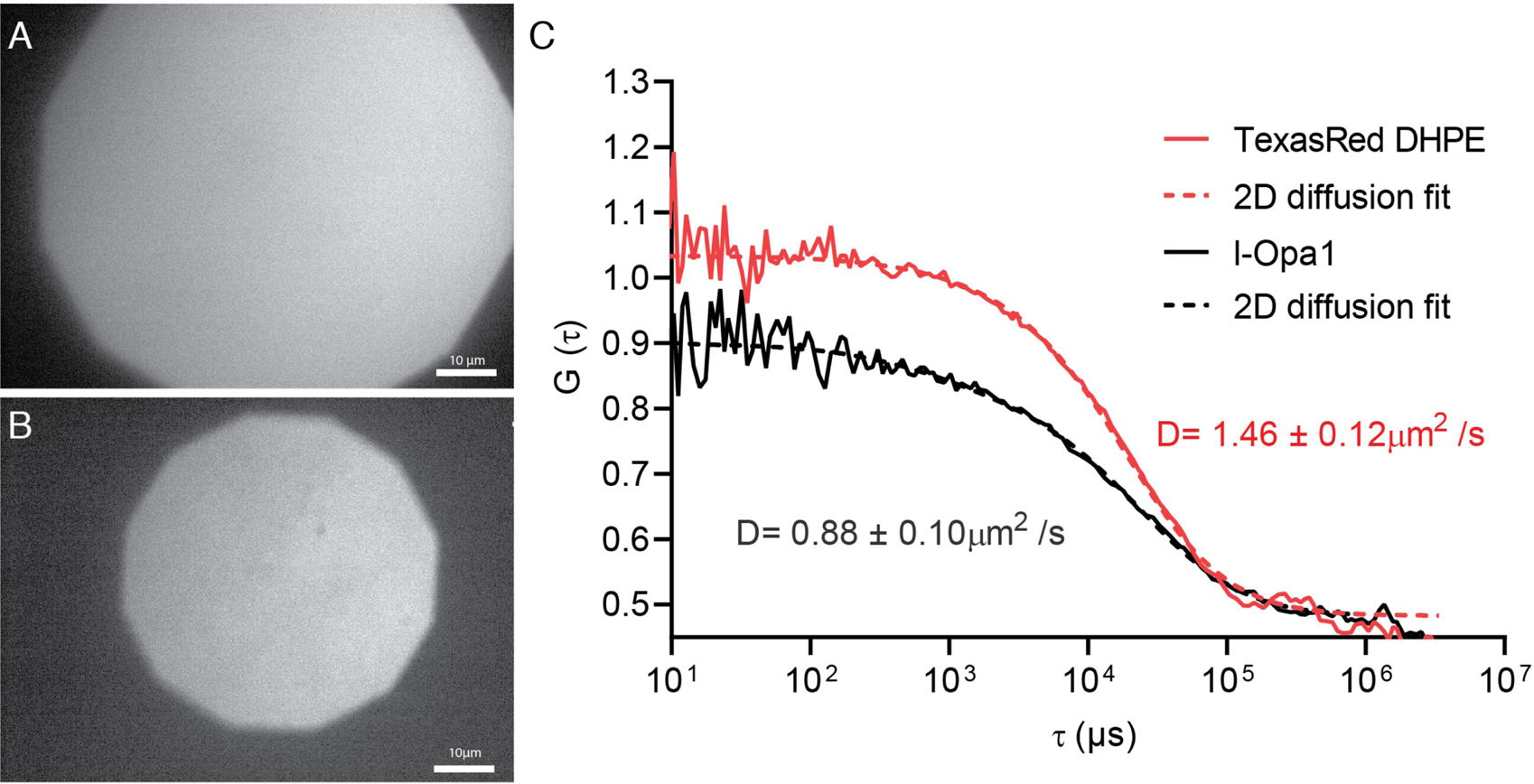
Epifluorescence image of polymer-tethered lipid bilayers before (A) and after Opa1 reconstitution (B), showing a homogeneous lipid bilayer. Scale bar: 10 µm. FCS profiles of TexasRed-PE and TexasRed labeled anti-Opa1 antibody show slower diffusion for reconstituted l-Opa1 (C), indicating successful reconstitution, and that the reconstituted l-Opa1 diffuses freely. Source data: Figure 2-fig sup 3-source data1.zip

**Figure 2 – figure supplement 4.**
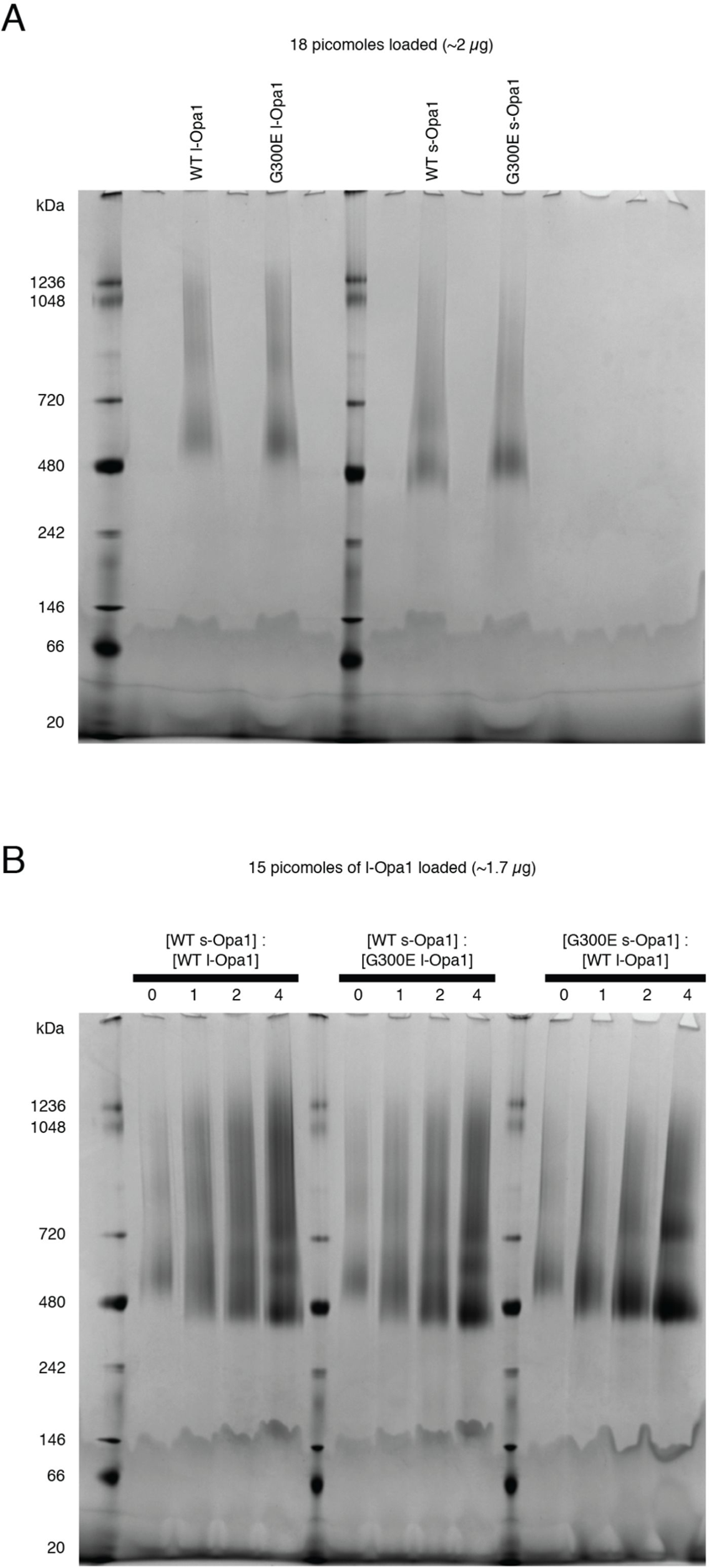
A. Blue native (BN-PAGE) gels show WT l-Opa1 and s-Opa1 can self-assemble as oligomers in DDM. B. Mixtures of WT l-Opa1 and WT s-Opa1 show a range of species from ∼480 KDa - ∼1 MDa. G300E l-Opa1, in the presence of WT s-Opa1, does not alter this gel migration pattern. In contrast, complexes comprising WT l-Opa1 and G300E s-Opa1 show a slight shift to a population mainly containing a ∼480 Kda and 720 KDa species.

**Figure 2 – figure supplement 5.**
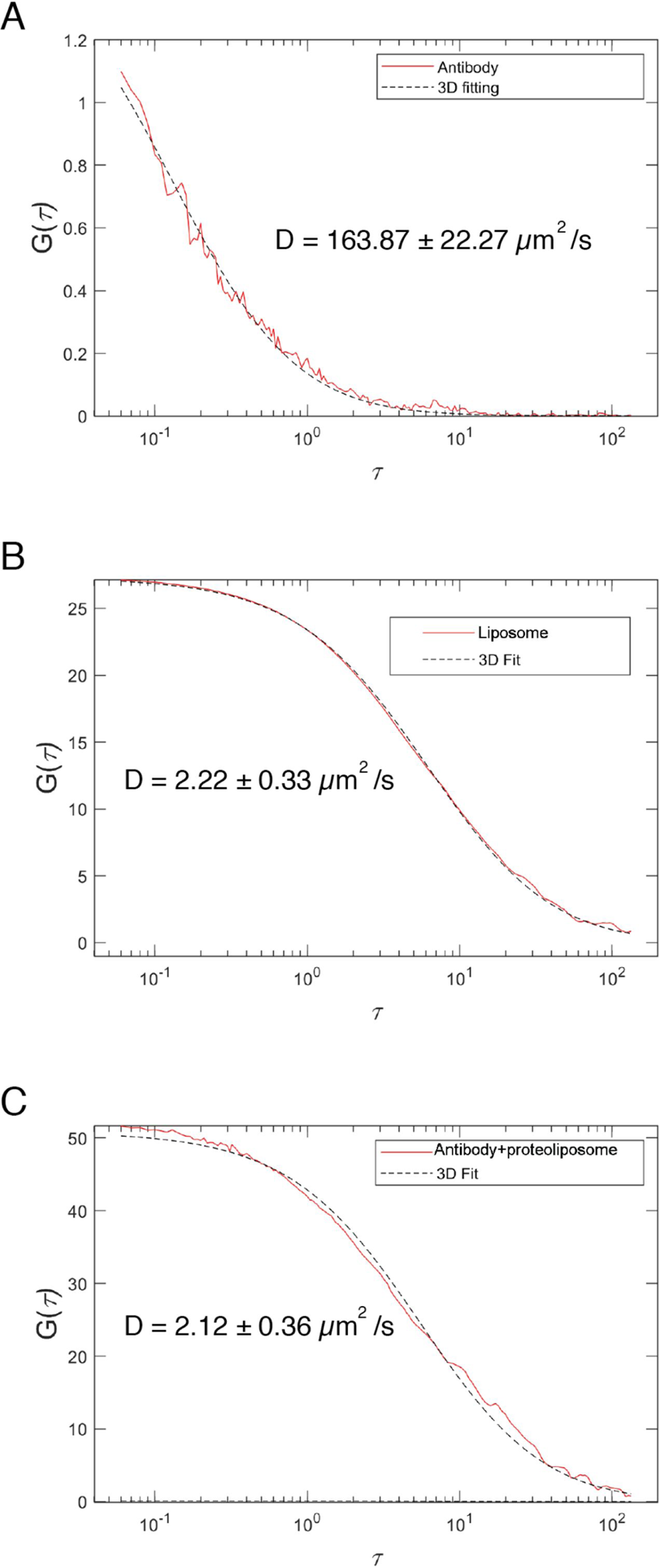
Fluorescence autocorrelation profiles of TexasRed labeled anti-His antibody in the presence of unlabeled liposomes (A), and TexasRed-PE-labeled liposomes (B), showing diffusion coefficients of unbound antibody versus liposomes. FCS profile of reconstituted l-Opa1 (detected with a TexasRed labeled antibody) (C) is similar to that of dye-labeled liposomes (B), indicating successful reconstitution of Opa1. Source data: Figure 2-fig sup 5-source data1.zip

**Figure 3 – figure supplement 1.**
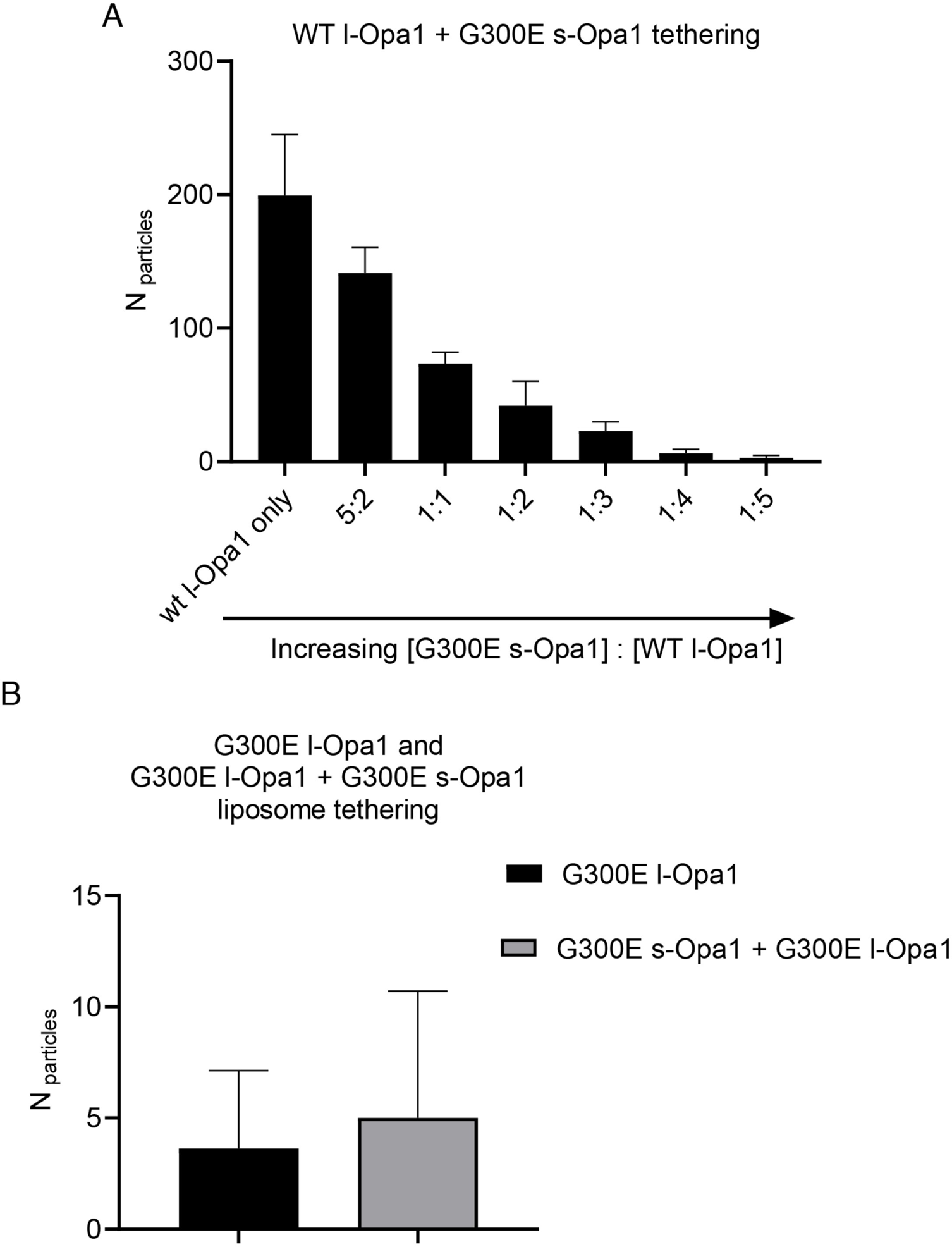
Effect of s-Opa1 competition on membrane tethering. Addition of G300E s-Opa1 detaches the l-Opa1 proteoliposomes tethered to l-Opa1-containing supported lipid (A). G300E l-Opa1 does not tether liposomes to a supported bilayer (B). G300E l-Opa1 in the presence of G300E s-Opa1 also does not tether membranes. Source data: Figure 3-fig sup 1-source data1.zip

**Figure 3 – figure supplement 2.**
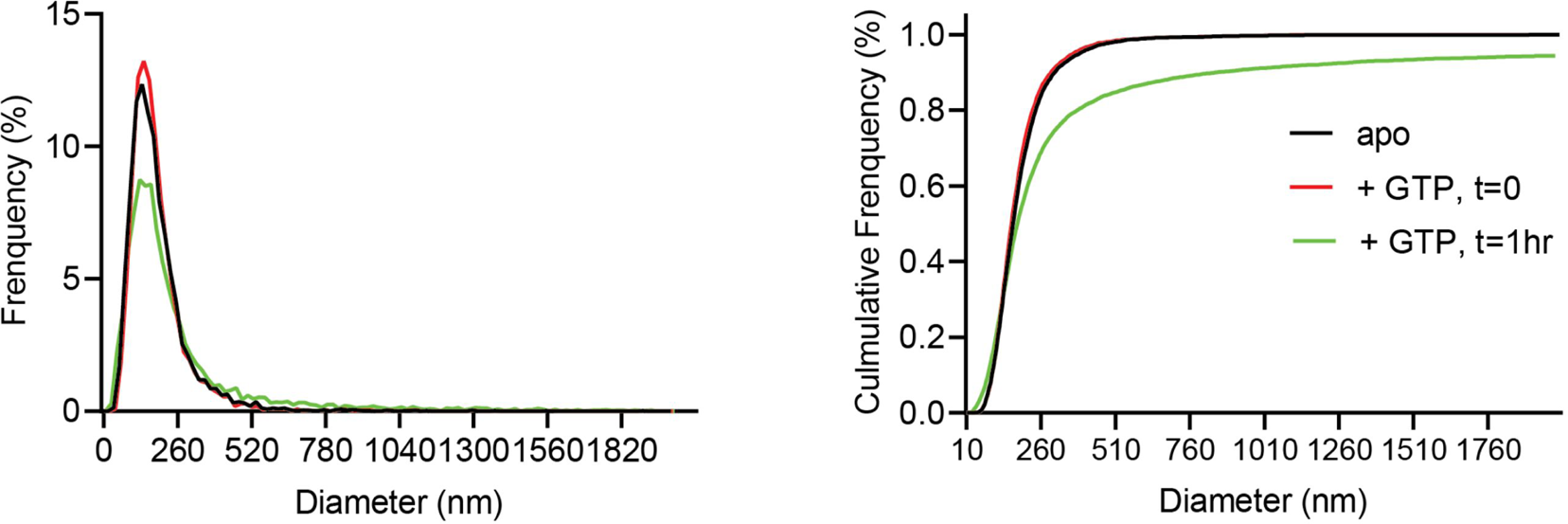
Normalized relative and cumulative size distributions show cardiolipin containing proteoliposomes shift to larger sizes 1 hour following GTP addition (green trace), as measured by Nanosight light scattering. Source data: Figure 3-fig sup 2-source data1.zip

**Figure 4 – figure supplement 1.**
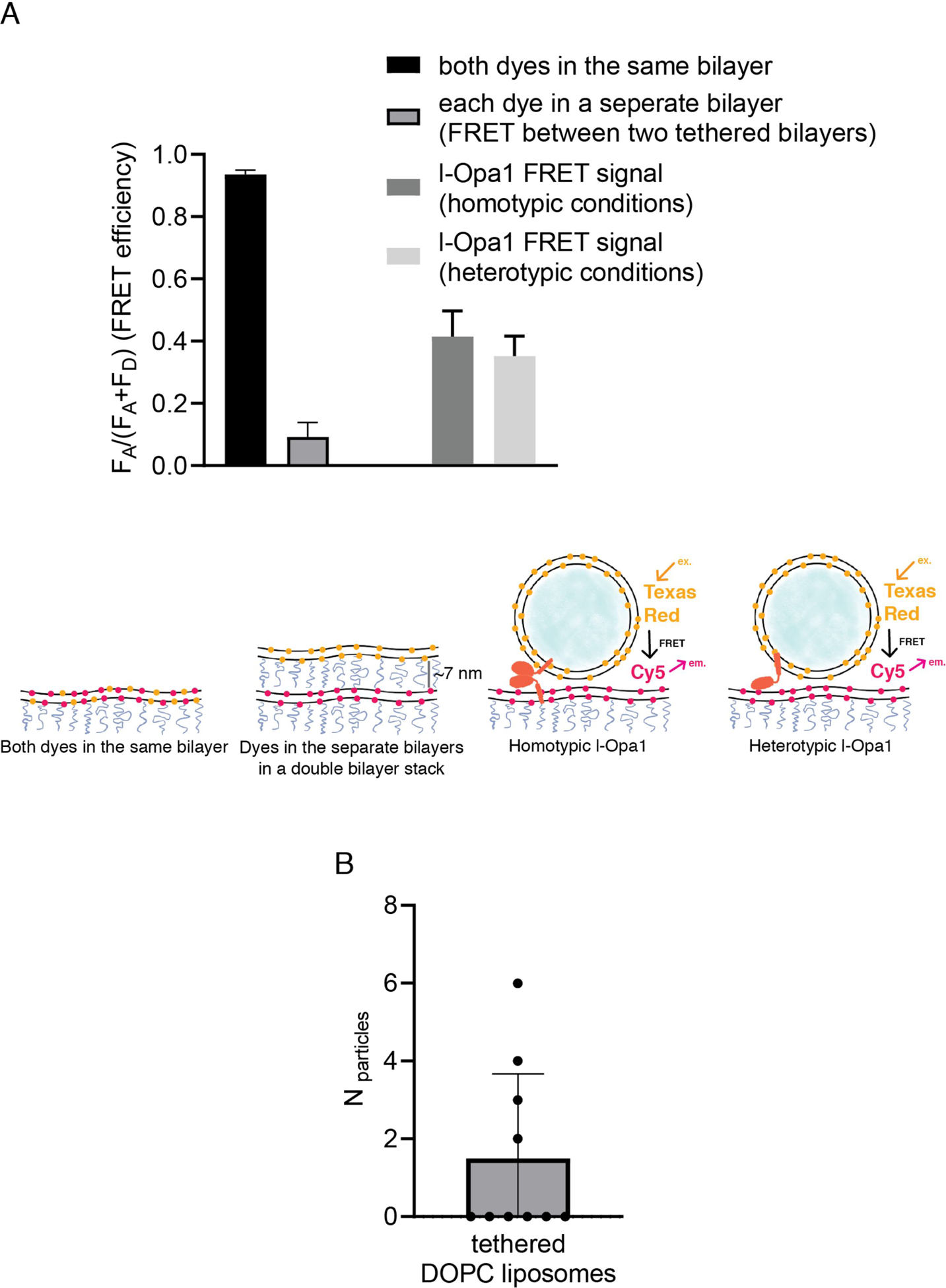
A. Controls for intra-membrane and inter-membrane FRET: When both TexasRed and Cy5 PE are present in the same bilayer, high FRET efficiency is observed. When TexasRed and Cy5 PE are present in two different bilayers, with a ∼7 nm tethering distance (from bilayer center to bilayer center in the double bilayer stack), FRET efficiency was low (data analyzed from 10 random spots in 2 bilayers (P<0.0001, t test). Analysis of ∼20 particles show ∼40% FRET efficiency for both homotypic and heterotypic tethering. This indicates that l-Opa1 is able to bring the two membranes within close proximity (< 7 nm) without mixing the two membranes. B. Quantification of DOPC liposomes tethered to a DOPC bilayer containing reconstituted l-Opa1. Liposomes do not tether to the supported bilayer, indicating that in the absence of cardiolipin, l-Opa1 does not tether liposomes alone. The lack of liposome docking to exposed regions also argues that few defects were introduced into the bilayer following reconstitution. Data from 3 different bilayers. Source data: Figure 4-fig sup 1-source data1.zip

**Figure 5 – figure supplement 1.**
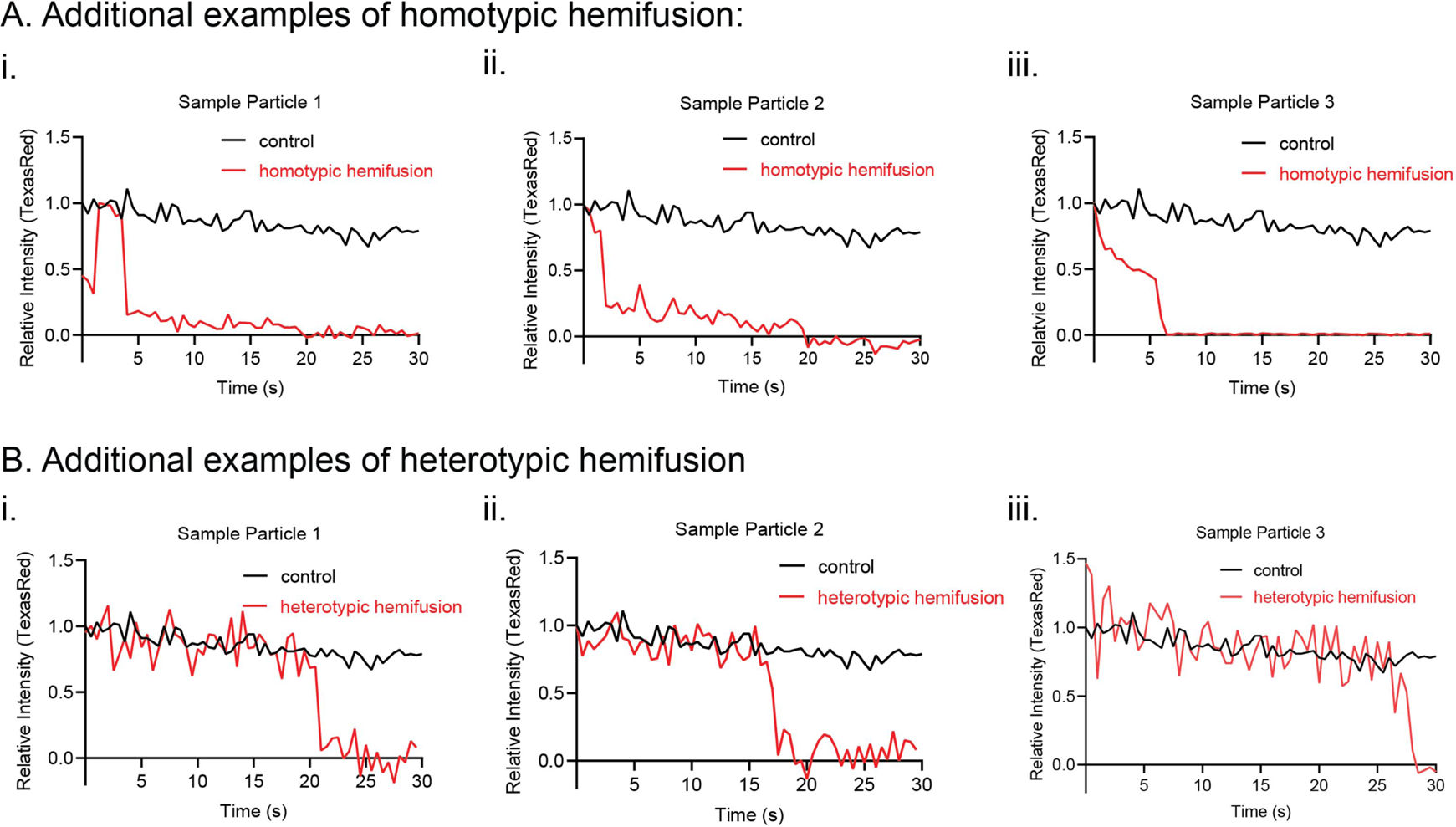
Additional kinetic traces for hemifusion curves under homotypic (A) and heterotypic (B) Opa1 hemifusion conditions. Control particle trace shown in black. Hemifusion trace shown in red. Source data: Figure 5-fig sup 1-source data1.zip

**Figure 6 – figure supplement 1.**
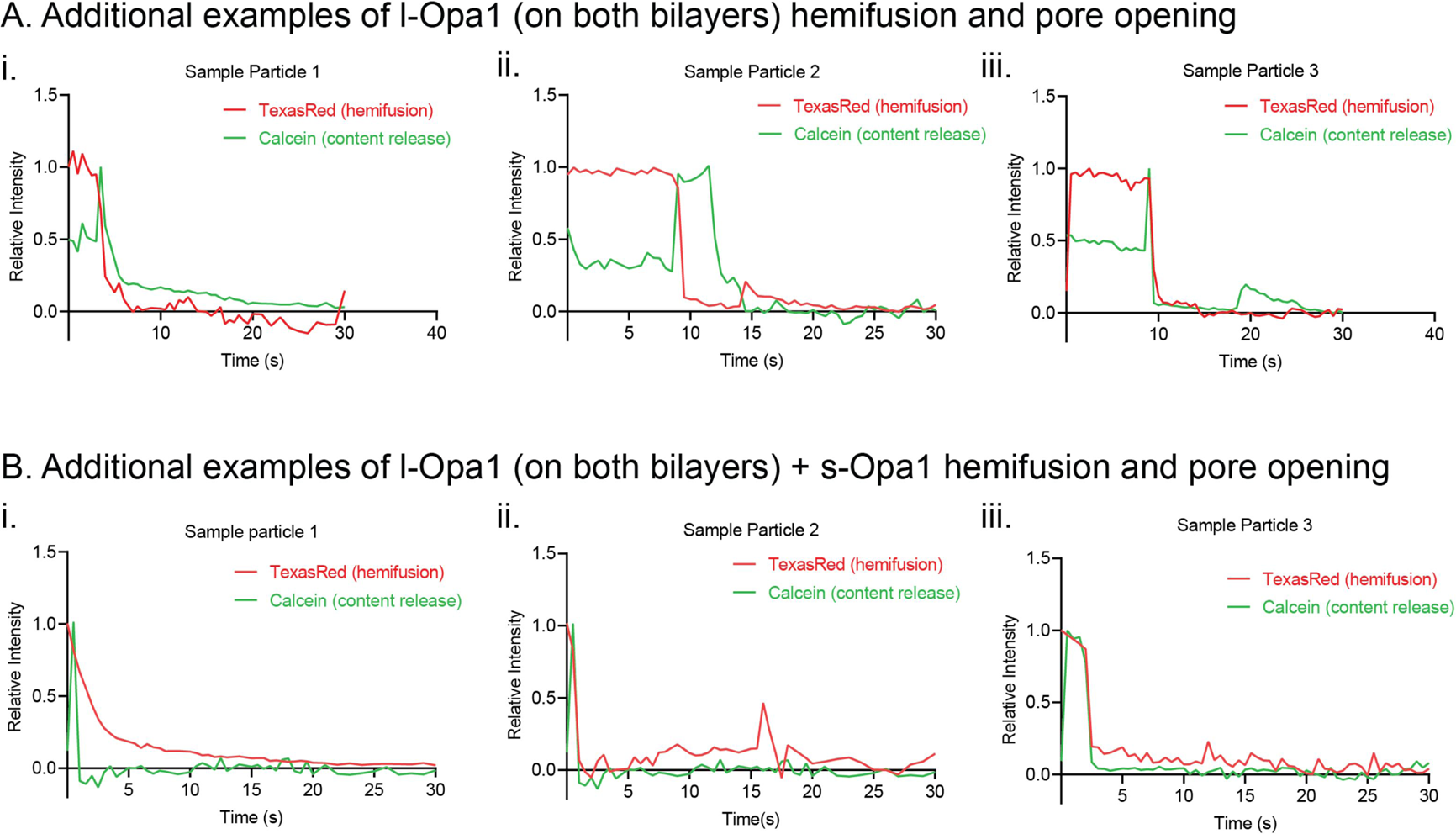
Additional kinetic traces for hemifusion and pore opening under homotypic l-Opa1 conditions (A), homotypic l-Opa1, and l-Opa1 + s-Opa1 (1:1) (B) conditions. Hemifusion (TexasRed) trace show in red. Pore opening (calcein, content release) trace shown in green. Figure 6-fig sup1-source data1.zip

## Notes

#### Summary of Updates

Revised manuscript

## References

1. Hoppins S, Lackner L, Nunnari J. The machines that divide and fuse mitochondria. Annu Rev Biochem. 2007;76:751–80. Epub 2007/03/17. doi: 10.1146/annurev.biochem.76.071905.090048. PubMed PMID: 17362197.

2. Cipolat S, Rudka T, Hartmann D, Costa V, Serneels L, Craessaerts K, Metzger K, Frezza C, Annaert W, D’Adamio L, Derks C, Dejaegere T, Pellegrini L, D’Hooge R, Scorrano L, De Strooper B. Mitochondrial rhomboid PARL regulates cytochrome c release during apoptosis via OPA1-dependent cristae remodeling. Cell. 2006;126(1):163–75. Epub 2006/07/15. doi: 10.1016/j.cell.2006.06.021. PubMed PMID: 16839884.

3. Cogliati S, Frezza C, Soriano ME, Varanita T, Quintana-Cabrera R, Corrado M, Cipolat S, Costa V, Casarin A, Gomes LC, Perales-Clemente E, Salviati L, Fernandez-Silva P, Enriquez JA, Scorrano L. Mitochondrial cristae shape determines respiratory chain supercomplexes assembly and respiratory efficiency. Cell. 2013;155(1):160–71. Epub 2013/09/24. doi: 10.1016/j.cell.2013.08.032. PubMed PMID: 24055366; PMCID: PMC3790458.

4. Nunnari J, Suomalainen A. Mitochondria: in sickness and in health. Cell. 2012;148(6):1145–59. doi: 10.1016/j.cell.2012.02.035. PubMed PMID: 22424226; PMCID: PMC5381524.

5. Westermann B. Mitochondrial fusion and fission in cell life and death. Nat Rev Mol Cell Biol. 2010;11(12):872–84. doi: 10.1038/nrm3013. PubMed PMID: 21102612.

6. Anand R, Wai T, Baker MJ, Kladt N, Schauss AC, Rugarli E, Langer T. The i-AAA protease YME1L and OMA1 cleave OPA1 to balance mitochondrial fusion and fission. J Cell Biol. 2014;204(6):919–29. doi: 10.1083/jcb.201308006. PubMed PMID: 24616225; PMCID: PMC3998800.

7. Chen H, Detmer SA, Ewald AJ, Griffin EE, Fraser SE, Chan DC. Mitofusins Mfn1 and Mfn2 coordinately regulate mitochondrial fusion and are essential for embryonic development. J Cell Biol. 2003;160(2):189–200. Epub 2003/01/16. doi: 10.1083/jcb.200211046. PubMed PMID: 12527753; PMCID: PMC2172648.

8. Alexander C, Votruba M, Pesch UE, Thiselton DL, Mayer S, Moore A, Rodriguez M, Kellner U, Leo-Kottler B, Auburger G, Bhattacharya SS, Wissinger B. OPA1, encoding a dynamin-related GTPase, is mutated in autosomal dominant optic atrophy linked to chromosome 3q28. Nat Genet. 2000;26(2):211–5. Epub 2000/10/04. doi: 10.1038/79944. PubMed PMID: 11017080.

9. Meeusen S, DeVay R, Block J, Cassidy-Stone A, Wayson S, McCaffery JM, Nunnari J. Mitochondrial inner-membrane fusion and crista maintenance requires the dynamin-related GTPase Mgm1. Cell. 2006;127(2):383–95. Epub 2006/10/24. doi: 10.1016/j.cell.2006.09.021. PubMed PMID: 17055438.

10. Meeusen S, McCaffery JM, Nunnari J. Mitochondrial fusion intermediates revealed in vitro. Science. 2004;305(5691):1747–52. Epub 2004/08/07. doi: 10.1126/science.1100612. PubMed PMID: 15297626.

11. MacVicar T, Langer T. OPA1 processing in cell death and disease - the long and short of it. J Cell Sci. 2016;129(12):2297–306. Epub 2016/05/18. doi: 10.1242/jcs.159186. PubMed PMID: 27189080.

12. Olichon A, Baricault L, Gas N, Guillou E, Valette A, Belenguer P, Lenaers G. Loss of OPA1 perturbates the mitochondrial inner membrane structure and integrity, leading to cytochrome c release and apoptosis. J Biol Chem. 2003;278(10):7743–6. Epub 2003/01/02. doi: 10.1074/jbc.C200677200. PubMed PMID: 12509422.

13. Pesch UE, Leo-Kottler B, Mayer S, Jurklies B, Kellner U, Apfelstedt-Sylla E, Zrenner E, Alexander C, Wissinger B. OPA1 mutations in patients with autosomal dominant optic atrophy and evidence for semi-dominant inheritance. Hum Mol Genet. 2001;10(13):1359–68. Epub 2001/07/07. doi: 10.1093/hmg/10.13.1359. PubMed PMID: 11440988.

14. Schmid SL, Frolov VA. Dynamin: functional design of a membrane fission catalyst. Annu Rev Cell Dev Biol. 2011;27:79–105. doi: 10.1146/annurev-cellbio-100109-104016. PubMed PMID: 21599493.

15. Ramachandran R, Schmid SL. The dynamin superfamily. Curr Biol. 2018; 28(8):R411-R6. doi: 10.1016/j.cub.2017.12.013. PubMed PMID: 29689225.

16. Faelber K, Dietrich L, Noel JK, Wollweber F, Pfitzner AK, Muhleip A, Sanchez R, Kudryashev M, Chiaruttini N, Lilie H, Schlegel J, Rosenbaum E, Hessenberger M, Matthaeus C, Kunz S, von der Malsburg A, Noe F, Roux A, van der Laan M, Kuhlbrandt W, Daumke O. Structure and assembly of the mitochondrial membrane remodelling GTPase Mgm1. Nature. 2019. doi: 10.1038/s41586-019-1372-3. PubMed PMID: 31292547.

17. Faelber K, Posor Y, Gao S, Held M, Roske Y, Schulze D, Haucke V, Noe F, Daumke O. Crystal structure of nucleotide-free dynamin. Nature. 2011;477(7366):556–60. Epub 2011/09/20. doi: 10.1038/nature10369. PubMed PMID: 21927000.

18. Mishra P, Carelli V, Manfredi G, Chan DC. Proteolytic cleavage of Opa1 stimulates mitochondrial inner membrane fusion and couples fusion to oxidative phosphorylation. Cell Metab. 2014;19(4):630–41. doi: 10.1016/j.cmet.2014.03.011. PubMed PMID: 24703695; PMCID: PMC4018240.

19. Ehses S, Raschke I, Mancuso G, Bernacchia A, Geimer S, Tondera D, Martinou JC, Westermann B, Rugarli EI, Langer T. Regulation of OPA1 processing and mitochondrial fusion by m-AAA protease isoenzymes and OMA1. J Cell Biol. 2009;187(7):1023–36. Epub 2009/12/30. doi: 10.1083/jcb.200906084. PubMed PMID: 20038678; PMCID: PMC2806285.

20. Ban T, Heymann JA, Song Z, Hinshaw JE, Chan DC. OPA1 disease alleles causing dominant optic atrophy have defects in cardiolipin-stimulated GTP hydrolysis and membrane tubulation. Hum Mol Genet. 2010;19(11):2113–22. doi: 10.1093/hmg/ddq088. PubMed PMID: 20185555; PMCID: PMC2865371.

21. Frezza C, Cipolat S, Martins de Brito O, Micaroni M, Beznoussenko GV, Rudka T, Bartoli D, Polishuck RS, Danial NN, De Strooper B, Scorrano L. OPA1 controls apoptotic cristae remodeling independently from mitochondrial fusion. Cell. 2006;126(1):177–89. Epub 2006/07/15. doi: 10.1016/j.cell.2006.06.025. PubMed PMID: 16839885.

22. Ford MG, Jenni S, Nunnari J. The crystal structure of dynamin. Nature. 2011;477(7366):561–6. Epub 2011/09/20. doi: 10.1038/nature10441. PubMed PMID: 21927001; PMCID: PMC4075756.

23. Antonny B, Burd C, De Camilli P, Chen E, Daumke O, Faelber K, Ford M, Frolov VA, Frost A, Hinshaw JE, Kirchhausen T, Kozlov MM, Lenz M, Low HH, McMahon H, Merrifield C, Pollard TD, Robinson PJ, Roux A, Schmid S. Membrane fission by dynamin: what we know and what we need to know. EMBO J. 2016;35(21):2270–84. Epub 2016/11/04. doi: 10.15252/embj.201694613. PubMed PMID: 27670760; PMCID: PMC5090216.

24. Chappie JS, Acharya S, Leonard M, Schmid SL, Dyda F. G domain dimerization controls dynamin’s assembly-stimulated GTPase activity. Nature. 2010;465(7297):435–40. Epub 2010/04/30. doi: 10.1038/nature09032. PubMed PMID: 20428113; PMCID: PMC2879890.

25. DeVay RM, Dominguez-Ramirez L, Lackner LL, Hoppins S, Stahlberg H, Nunnari J. Coassembly of Mgm1 isoforms requires cardiolipin and mediates mitochondrial inner membrane fusion. J Cell Biol. 2009;186(6):793–803. doi: 10.1083/jcb.200906098. PubMed PMID: 19752025; PMCID: PMC2753158.

26. Herlan M, Vogel F, Bornhovd C, Neupert W, Reichert AS. Processing of Mgm1 by the rhomboid-type protease Pcp1 is required for maintenance of mitochondrial morphology and of mitochondrial DNA. J Biol Chem. 2003;278(30):27781–8. Epub 2003/04/23. doi: 10.1074/jbc.M211311200. PubMed PMID: 12707284.

27. Song Z, Chen H, Fiket M, Alexander C, Chan DC. OPA1 processing controls mitochondrial fusion and is regulated by mRNA splicing, membrane potential, and Yme1L. J Cell Biol. 2007;178(5):749–55. Epub 2007/08/22. doi: 10.1083/jcb.200704110. PubMed PMID: 17709429; PMCID: PMC2064540.

28. Ban T, Ishihara T, Kohno H, Saita S, Ichimura A, Maenaka K, Oka T, Mihara K, Ishihara N. Molecular basis of selective mitochondrial fusion by heterotypic action between OPA1 and cardiolipin. Nat Cell Biol. 2017;19(7):856–63. doi: 10.1038/ncb3560. PubMed PMID: 28628083.

29. Del Dotto V, Fogazza M, Carelli V, Rugolo M, Zanna C. Eight human OPA1 isoforms, long and short: What are they for? Biochim Biophys Acta Bioenerg. 2018;1859(4):263–9. Epub 2018/02/01. doi: 10.1016/j.bbabio.2018.01.005. PubMed PMID: 29382469.

30. Naumann C, Prucker O, Lehmann T, Ruhe J, Knoll W, Frank CW. The polymer-supported phospholipid bilayer: tethering as a new approach to substrate-membrane stabilization. Biomacromolecules. 2002;3(1):27–35. Epub 2002/02/28. PubMed PMID: 11866552.

31. Siegel AP, Kimble-Hill A, Garg S, Jordan R, Naumann CA. Native ligands change integrin sequestering but not oligomerization in raft-mimicking lipid mixtures. Biophys J. 2011;101(7):1642–50. Epub 2011/10/04. doi: 10.1016/j.bpj.2011.08.040. PubMed PMID: 21961590; PMCID: PMC3183796.

32. Liu TY, Bian X, Romano FB, Shemesh T, Rapoport TA, Hu J. Cis and trans interactions between atlastin molecules during membrane fusion. Proc Natl Acad Sci U S A. 2015;112(15):E1851–60. Epub 2015/04/01. doi: 10.1073/pnas.1504368112. PubMed PMID: 25825753; PMCID: PMC4403200.

33. O’Donnell JP, Cooley RB, Kelly CM, Miller K, Andersen OS, Rusinova R, Sondermann H. Timing and Reset Mechanism of GTP Hydrolysis-Driven Conformational Changes of Atlastin. Structure. 2017;25(7):997–1010 e4. Epub 2017/06/13. doi: 10.1016/j.str.2017.05.007. PubMed PMID: 28602821; PMCID: PMC5516944.

34. Abutbul-Ionita I, Rujiviphat J, Nir I, McQuibban GA, Danino D. Membrane tethering and nucleotide-dependent conformational changes drive mitochondrial genome maintenance (Mgm1) protein-mediated membrane fusion. J Biol Chem. 2012;287(44):36634–8. Epub 2012/09/15. doi: 10.1074/jbc.C112.406769. PubMed PMID: 22977249; PMCID: PMC3481265.

35. Chao LH, Klein DE, Schmidt AG, Pena JM, Harrison SC. Sequential conformational rearrangements in flavivirus membrane fusion. Elife. 2014;3:e04389. Epub 2014/12/06. doi: 10.7554/eLife.04389. PubMed PMID: 25479384; PMCID: PMC4293572.

36. Ivanovic T, Choi JL, Whelan SP, van Oijen AM, Harrison SC. Influenza-virus membrane fusion by cooperative fold-back of stochastically induced hemagglutinin intermediates. Elife. 2013;2:e00333. Epub 2013/04/04. doi: 10.7554/eLife.00333. PubMed PMID: 23550179; PMCID: PMC3578201.

37. Rawle RJ, van Lengerich B, Chung M, Bendix PM, Boxer SG. Vesicle fusion observed by content transfer across a tethered lipid bilayer. Biophys J. 2011;101(8):L37–9. Epub 2011/10/19. doi: 10.1016/j.bpj.2011.09.023. PubMed PMID: 22004762; PMCID: PMC3192961.

38. Griparic L, Kanazawa T, van der Bliek AM. Regulation of the mitochondrial dynamin-like protein Opa1 by proteolytic cleavage. J Cell Biol. 2007;178(5):757–64. Epub 2007/08/22. doi: 10.1083/jcb.200704112. PubMed PMID: 17709430; PMCID: PMC2064541.

39. Lee H, Smith SB, Yoon Y. The short variant of the mitochondrial dynamin OPA1 maintains mitochondrial energetics and cristae structure. J Biol Chem. 2017;292(17):7115–30. doi: 10.1074/jbc.M116.762567. PubMed PMID: 28298442; PMCID: PMC5409478.

40. Baker MJ, Lampe PA, Stojanovski D, Korwitz A, Anand R, Tatsuta T, Langer T. Stress-induced OMA1 activation and autocatalytic turnover regulate OPA1-dependent mitochondrial dynamics. EMBO J. 2014;33(6):578–93. doi: 10.1002/embj.201386474. PubMed PMID: 24550258; PMCID: PMC3989652.

41. Brandt T, Cavellini L, Kuhlbrandt W, Cohen MM. A mitofusin-dependent docking ring complex triggers mitochondrial fusion in vitro. Elife. 2016;5. Epub 2016/06/03. doi: 10.7554/eLife.14618. PubMed PMID: 27253069; PMCID: PMC4929004.

42. Zick M, Duvezin-Caubet S, Schafer A, Vogel F, Neupert W, Reichert AS. Distinct roles of the two isoforms of the dynamin-like GTPase Mgm1 in mitochondrial fusion. FEBS Lett. 2009;583(13):2237–43. Epub 2009/06/10. doi: 10.1016/j.febslet.2009.05.053. PubMed PMID: 19505460.

43. Wai T, Langer T. Mitochondrial Dynamics and Metabolic Regulation. Trends Endocrinol Metab. 2016;27(2):105–17. doi: 10.1016/j.tem.2015.12.001. PubMed PMID: 26754340.

44. Duvezin-Caubet S, Jagasia R, Wagener J, Hofmann S, Trifunovic A, Hansson A, Chomyn A, Bauer MF, Attardi G, Larsson NG, Neupert W, Reichert AS. Proteolytic processing of OPA1 links mitochondrial dysfunction to alterations in mitochondrial morphology. J Biol Chem. 2006;281(49):37972–9. Epub 2006/09/28. doi: 10.1074/jbc.M606059200. PubMed PMID: 17003040.

45. Rainbolt TK, Lebeau J, Puchades C, Wiseman RL. Reciprocal Degradation of YME1L and OMA1 Adapts Mitochondrial Proteolytic Activity during Stress. Cell Rep. 2016;14(9):2041–9. doi: 10.1016/j.celrep.2016.02.011. PubMed PMID: 26923599; PMCID: PMC4785047.

46. Ishihara N, Fujita Y, Oka T, Mihara K. Regulation of mitochondrial morphology through proteolytic cleavage of OPA1. EMBO J. 2006;25(13):2966–77. Epub 2006/06/17. doi: 10.1038/sj.emboj.7601184. PubMed PMID: 16778770; PMCID: PMC1500981.

47. Baker MJ, Tatsuta T, Langer T. Quality control of mitochondrial proteostasis. Cold Spring Harb Perspect Biol. 2011;3(7). doi: 10.1101/cshperspect.a007559. PubMed PMID: 21628427; PMCID: PMC3119916.

48. Zhang D, Zhang Y, Ma J, Niu T, W. C, X. P, Zhia Y, Sun F. Cryo-EM structures reveal interactions of S-OPA1 with membrane and changes upon nucleotide binding. bioRxiv. 2019. doi: https://doi.org/10.1101/528042.

49. Ge Y, Siegel AP, Jordan R, Naumann CA. Ligand binding alters dimerization and sequestering of urokinase receptors in raft-mimicking lipid mixtures. Biophys J. 2014;107(9):2101–11. Epub 2014/11/25. doi: 10.1016/j.bpj.2014.09.021. PubMed PMID: 25418095; PMCID: PMC4223190.

50. Lewis RN, McElhaney RN. The physicochemical properties of cardiolipin bilayers and cardiolipin-containing lipid membranes. Biochim Biophys Acta. 2009;1788(10):2069–79. Epub 2009/03/31. doi: 10.1016/j.bbamem.2009.03.014. PubMed PMID: 19328771.

51. Minner DE, Herring VL, Siegel AP, Kimble-Hill A, Johnson MA, Naumann CA. Iterative layer-by-layer assembly of polymer-tethered multi-bilayers using maleimide-thiol coupling chemistry. Soft Matter. 2013;9(40):9643–50. Epub 2013/10/28. doi: 10.1039/c3sm51446c. PubMed PMID: 26029773.

52. Huang Y, Bharill S, Karandur D, Peterson SM, Marita M, Shi X, Kaliszewski MJ, Smith AW, Isacoff EY, Kuriyan J. Molecular basis for multimerization in the activation of the epidermal growth factor receptor. Elife. 2016;5. Epub 2016/03/29. doi: 10.7554/eLife.14107. PubMed PMID: 27017828; PMCID: PMC4902571.

53. Comar WD, Schubert SM, Jastrzebska B, Palczewski K, Smith AW. Time-resolved fluorescence spectroscopy measures clustering and mobility of a G protein-coupled receptor opsin in live cell membranes. J Am Chem Soc. 2014;136(23):8342–9. doi: 10.1021/ja501948w. PubMed PMID: 24831851; PMCID: PMC4063175.

54. Jaqaman K, Loerke D, Mettlen M, Kuwata H, Grinstein S, Schmid SL, Danuser G. Robust single-particle tracking in live-cell time-lapse sequences. Nat Methods. 2008;5(8):695–702. Epub 2008/07/22. doi: 10.1038/nmeth.1237. PubMed PMID: 18641657; PMCID: PMC2747604.

